# Decoding stress specific transcriptional regulation by causality aware Graph-Transformer deep learning

**DOI:** 10.1101/2025.03.11.642297

**Authors:** Umesh Bhati, Akanksha Sharma, Sagar Gupta, Anchit Kumar, Upendra Kumar Pradhan, Ravi Shankar

**Author notes:** **Authors’ email addresses:** UB AS SG AK UKP RS.

## Abstract

Cells respond to environmental stimuli through transcriptional reprogramming orchestrated by transcription factors (TFs), which interpret cis-regulatory DNA sequences to determine the timing and location of gene expression. The diversification of TFs and their interactions with cis-regulatory elements (CREs) underpins plant adaptation to stress through the formation of gene regulatory networks (GRNs). However, deciphering condition-specific GRNs and identifying transcription factor binding motifs (TFBMs) for spatio-temporal gene expression remain major challenges in plant biology. To decipher the conditional networks governing TF-Target gene interactions, we developed CTF-BIND, a novel computational framework designed to reason about the spatio-temporal dynamics of TF activity. Leveraging over ∼23TB of multi-omics data (ChIP-seq, RNA-seq, and protein-protein interaction data), we constructed Bayesian causal networks capable of explaining TF activity across diverse conditions. These networks, validated against extensive experimental data, were then integrated into a Graph Transformer deep learning system. This system uses expression information of network components to quantitatively determine TF activity levels. Models were developed for 110 abiotic stress-related TFs, enabling accurate condition-specific detection of TF binding directly from RNA-seq data, eliminating the need for separate ChIP-seq experiments. CTF-BIND achieved a high average accuracy of ∼93% when tested against experimentally established data from various conditions. It is implemented as an interactive, open-access web server, it not only provides TF binding profiles but also facilitates downstream functional analysis. Furthermore, we developed CTF-BIND-DB, (https://hichicob.ihbt.res.in/ctfbind/) a database capturing dynamic shifts in regulatory pathways, providing information on TGs, network ontology, and binding motifs. CTF-BIND and CTF-BIND-DB represent a transformative approach for understanding and determining TF activity in plant stress responses, offering a powerful tool for crop improvement and bypassing the limitations of traditional methods and extensive experimental validation.

## Introduction

Plants are continually exposed to diverse environmental stressors in their natural habitats, survive through sophisticated adaptive mechanisms (Zhu et al. 2016). The activation of plant stress responses involves a complex, multi-dimensional process characterized by large-scale transcriptional reprogramming, requiring an intricate regulatory system (Kapoor et al. 2021; Kim et al. 2021; Chae et al. 2012). Gene expression variations in response to environmental stimuli contribute to phenotypic diversity within species, leading to adaptive evolution (Hämälä et al. 2022; Pimpinelli et al. 2020; Garfield et al. 2013). Plants exhibit both stress-specific adaptive responses and global protective mechanisms against multiple environmental stressors. These responses involve multiple stress perception and signaling pathways, some of which are stress-specific, while others demonstrate cross-talk at various stages of the stress response process (Boone et al. 2023; López-Maury et al. 2017). Despite extensive research on plant responses to individual stress conditions across various model organisms (Meng et al. 2020; Zhang et al. 2017; Naika et al. 2013), the underlying differentiation of transcriptional regulatory networks among diverse abiotic conditions within a given plant species remains largely unexplored.

Transcriptional regulation, a fundamental process governing gene expression and orchestrating cellular activities, plays a crucial role in shaping phenotypic diversity. This process is driven by transcription factors (TFs), which modulate the expression of their target genes (TGs) by binding to specific DNA sequences known as TF binding sites (TFBSs). The entirety of regulatory interactions between TFs and their TGs constitutes a gene regulatory network (GRN), providing a systems-level view of transcriptional control. While TFs are often categorized as activators or repressors, gene regulation is typically a complex process involving combinatorial control by multiple TFs, with the specific context (e.g., developmental stage, environmental conditions) dictating how a TF influence TGs expression (Martinez-Corral et al., 2024; Li et al., 2015). The *Arabidopsis thaliana* genome, for instance, encodes approximately 2,000 TFs (Wang et al., 2021; Weirauch et al., 2014); however, experimentally validated regulatory interactions are available for a considerably smaller subset (fewer than 250) (Davuluri et al., 2003; de Velde et al., 2014), highlighting the need for robust methods to decipher GRNs. When plants encounter stress, specific TFs are activated, initiating a rapid signal transduction cascade that affects the expression of numerous downstream genes (Zhu et al., 2016). This transcriptional reprogramming is crucial for mounting appropriate adaptive responses **(Figure 1)**.

**Figure 1:**
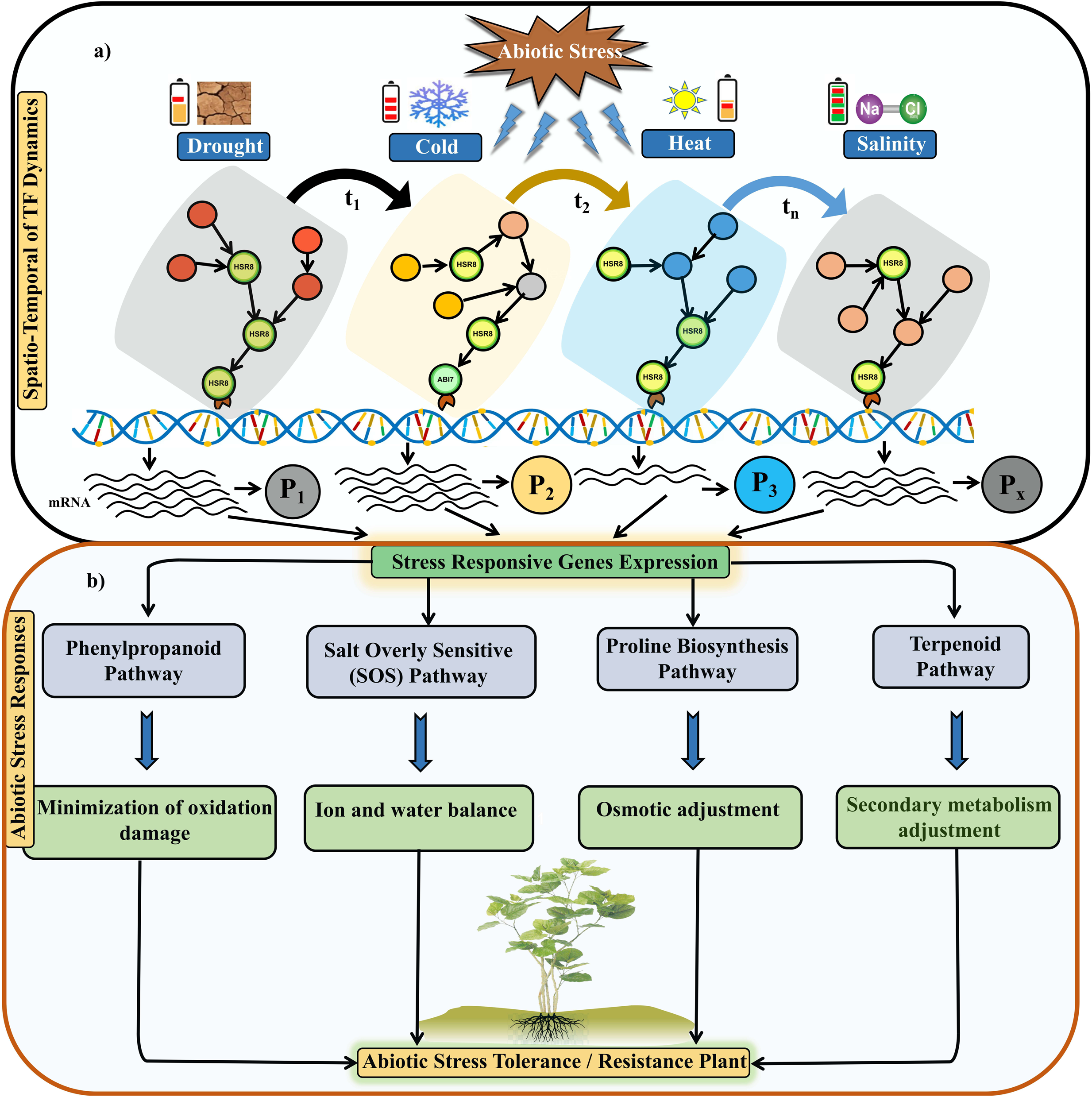
**(a) Spatio-Temporal Dynamics of TF Activities under Abiotic Stress Conditions**. This figure illustrates the spatio-temporal dynamics of TFs in response to various abiotic stress conditions such as drought, cold, heat, and salinity. The figure shows distinct time points (t_₁_, t_₂_, t_ₙ_) during which different sets of TFs (represented by colored circles) become activated. Each stress condition triggers a unique set of TFs, which bind to specific promoter regions of target genes, resulting in the transcription of stress-responsive genes. The progression from t₁ to tₙ highlights the sequential and dynamic nature of TF activation, indicating how plants orchestrate a coordinated response to cope with stress. **b) Pathways Involved in Abiotic Stress Responses and Their Impact on Plant Tolerance** This depicts the downstream effects of stress-responsive gene expression on various physiological pathways. Stress-induced TF activities lead to the activation of multiple pathways. These pathways collectively enhance the plant’s tolerance and resistance to abiotic stress by enabling various adaptive mechanisms such as antioxidation, ion homeostasis, osmotic regulation, and metabolic adjustments. The integration of these responses culminates in an overall increase in abiotic stress tolerance, as depicted by the resilient plant at the center.

Consequently, significant efforts have been devoted to developing computational algorithms for inferring GRNs from gene expression data, identifying regulatory modules, and determining condition-specific regulators (Velde et al., 2014; Segal et al., 2003; Babu et al., 2009). Network-based approaches have proven particularly effective in capturing the inherent complexity of biological systems (Bassel et al., 2022; Tuteja et al., 2019). From a network perspective, TF-TG interactions can be represented as directed graphs, depicting transcriptional modulation (Alvarez et al., 2011), often exhibiting scale-free topology with highly connected regulatory hubs (Barabasi et al., 2004).

Studies have begun to unravel GRNs in specific cellular contexts using various experimental techniques. For instance, Brady et al. (2011) utilized a gene-centric yeast one-hybrid (Y1H) approach to elucidate the first tissue-specific GRN in *Arabidopsis* roots under different abiotic stress conditions. Similarly, Taylor-Teeples (2015) employed Y1H to investigate the GRN underlying secondary cell wall synthesis. In contrast to in vitro methods like Y1H, *in vivo* techniques such as chromatin immunoprecipitation followed by sequencing (ChIP-seq) allow for the direct monitoring of TF binding to genomic DNA, thereby mapping the targets of individual TFs (Kaufmann et al., 2010; Xie et al., 2017). Additionally, open chromatin profiling integrates TF footprint information obtained from DNase I hypersensitivity assays with known TFBSs to construct unbiased GRNs, as it does not require prior knowledge of potential TFs or target genes (Friedel et al., 2022).

Despite these advancements, a comprehensive systems-level understanding of GRNs that orchestrate the complex abiotic stress response remains elusive (Collin et al. 2021; Cramer et al., 2011). While coexpression analysis through statistical correlation and clustering can infer functional associations based on the guilt-by-association principle, the specific regulators and dynamics of regulation remain unclear. Resources such as Genevestigator (Zimmermann et al., 2004), ATTED-II (Obayashi et al., 2022), and NetREx (Sircar et al., 2022) have facilitated the generation of co-expression-based gene association networks for *Arabidopsis thaliana*, yet the detailed regulatory interactions between TFs and their target genes have only been mapped for a limited number of interactions (Brady et al., 2021; Gaudinier et al., 2011) and largely miss to present the reasoning circuit behind them.

This lack of comprehensive, condition-specific regulatory information hinders our ability to understand the causal mechanisms driving plant stress responses. While methods like Y1H (Brady et al., 2021; Taylor-Teeples et al., 2015) and ChIP-seq (Kaufmann et al., 2010; Xie et al., 2010) provide valuable data on direct TF-TG interactions, they do not fully capture the dynamic regulation of TF activity across diverse conditions. The observed variability in TF interactions across different conditions highlights the need for a more holistic approach that considers the context-dependent nature of transcriptional regulation. Therefore, there is a critical need for methods that not only identify TF-TG interactions but also establish causal relationships and predict TF influence under specific environmental conditions.

Although some of these interactions have been functionally validated for a limited number of TF-TG pairs and conditions, experimentally verifying all ChIP-seq interactions across a broader spectrum of conditions remains challenging and costly affair. Moreover, previous studies has not fully integrated high-throughput data from diverse platforms such as RNA-seq and ChIP-seq to construct reasoning circuits that could explain TF activation through resilient models. These investigations have, however, emphasized the critical need to elucidate the spatio-temporal dynamics of TFs and strongly advocated for a more holistic approach to interaction analysis – a challenge our present study aims to address with respect to abiotic stress. Developing such reasoning frameworks is crucial for both explaining observed phenomena and establishing TF behavior across a wide range of conditions. In this context, computational modeling of these interactions emerges as an urgent necessity. Recently, our group had applied bayesian causal network analysis to detect RBPs-RNA interactions to computationally profile miRNA (Pradhan et al. 2021). A similar approach can be useful for TF-binding timing and associated responses.

To address the above mentioned challenged and limitations, we conducted a comprehensive analysis encompassing 46 abiotic stress conditions, leveraging over 23 TB of multi-omics data, including protein-DNA interactions (ChIP-seq), protein-protein interaction (PPI), and transcriptome profiling (RNA-seq) for *A. thaliana*. This integrative approach enabled us to achieve three key objectives:

**(1)** Construction of causal networks and logical units to elucidate the spatio-temporal dynamics of TF activity and its impact on target gene expression, employing Bayesian network analysis to decipher condition-specific causal networks.
**(2)** These Bayesian networks were then used to train a deep learning model, a Graph Transformer termed CTF-BIND, capable of accurately identifying TF binding under diverse conditions, achieving over ∼93% accuracy for a model system encompassing 110 Tfs.
**(3)** Integrating the insights from the Bayesian networks and CTF-BIND, we developed CTF-BIND-DB, a novel database for 110 abiotic stress-responsive TFs. CTF-BIND-DB uniquely captures the dynamic shifts in regulatory pathways across conditions, providing information on target genes, network ontology, and binding spots. This database is integrated with a user-friendly web server, allowing researchers to identify spatio-temporal TF binding to user-provided sequences. To our knowledge, CTF-BIND-DB is the first of its kind resource to provide such comprehensive condition-specific information on TF binding dynamics, in general as well as abiotic stress biology.

This study present, causal networks of TFs across various abiotic stress conditions, elucidating both stress-specific and global regulatory pathways in relation to stress tolerance. A key outcome of this work is the development of CTF-BIND, a system that significantly advancement by enabling accurate identification of condition-specific TF binding profiles without requiring new ChIP-seq experiments. By leveraging the dynamic information embedded within the learned TF-TG Bayesian networks and expression, CTF-BIND offers unprecedented accuracy. Rigorous testing against experimentally validated data from diverse conditions has demonstrated its high accuracy and reliability. This integrative approach, combining large-scale genomic and advanced computational modeling, bypasses the limitations of traditional TFBS mapping and the need for extensive experimental validation. CTF-BIND streamlines the identification of key regulatory interactions under various abiotic stresses and provides a powerful tool for developing targeted genetic modification strategies to enhance plant adaptation to environmental stressors. The study opens a new gateway to further extend its methodology for condition specific TF binding and gene regulation, which could answer and provide solutions for several challenges in plant biology, the resources presented here are bound to be invaluable ones.

## Materials and methods

### Data collection and processing of gene expression data

A comprehensive dataset of 780 gene expression samples (RNA-seq) representing 46 conditions for *A. thaliana* across the four major abiotic stress types drought, heat, cold, and salinity were curated from the European Nucleotide Archive (ENA) and the AtGenExpress platform. 278 TFs associated with abiotic stress regulation were retrieved from the STIFDB2 (Stress Responsive Transcription Factor Database) (Naika et al., 2012), focusing on regulatory elements critical for stress response. For this analysis, time-series gene expression data for cold, heat, drought, and salinity stress were extracted from shoot and root tissues of *A. thaliana*, with raw data obtained from the GEO database (Barrett et al., 2013). To isolate the immediate stress response, stress recovery phase data were excluded, and differential expression profiles (DEGs) were generated. The dataset was organized into cold stress (180 samples, 7 time points), heat stress (221 samples, 9 time points), salt stress (277 samples, 5 time points), and drought stress (102 samples, 7 time points), providing a detailed, time-resolved view of transcriptional dynamics under abiotic stress. All details of samples summarized in **Supplementary Table 1**.

The gene expression data (RNA-seq fastq file) was filtered using filteR while using their quality guideline (Gahlan et at. 2012). For mapping of these RNA-seq reads across the *Arabidopsis* genome, Hisat2 was used which is based on Bowtie platform (Langmead et al. 2012). The alignment results were saved in SAM format for expression quantification. Ht-count was used for quantification of gene expression from SAM files. Furthermore, the quantification counts were normalized to fragments per kilo base per million mapped reads (FPKM) for paired end reads and reads per kilo base per million mapped reads (RPKM) for single end reads, using in-house developed shell scripts for every single gene across different conditions. To address technical variability, batch effect correction was performed using the limma R-package (Ritchie et al., 2015).

### ChIP-seq data collection and processing

490 ChIP-seq samples representing 124 TFs for *A. thaliana* were retrieved from public repositories, including ChIP-Hub (https://www.chip-hub.org/), PlantPan 3.0 (http://plantpan.itps.ncku.edu.tw/), and the Gene Expression Omnibus (GEO) (https://www.ncbi.nlm.nih.gov/geo/) (**Supplementary Table 2**). This initial dataset comprised approximately 9.8 TB of raw sequencing reads. To ensure high data quality, stringent quality control and filtering procedures were implemented. Samples exhibiting low sequencing depth, poor signal-to-noise ratios, or other quality issues were excluded from further analysis. We applied a peak filtering step, removing samples where the total number of identified peaks was less than 400. This threshold was chosen to prioritize samples with sufficient binding signal to confidently identify TF-TG interactions. This rigorous filtering resulted in a refined dataset encompassing 110 TFs, which were subsequently considered for downstream analysis related to abiotic stress responses.

### Reconstruction of TF:TG associations using Bayesian network analysis (BNA)

ChIP-seq and RNA-seq data having same experimental conditions in *A. thaliana* were collected from ChIP-Hub, GEO and ENA databases. Total 22 experimental conditions were common among both the two types of high-throughput data. Possible interacting partners were collected for each TF from various resources like STRING, AGRIS and stress knowledge map database (SKM) (Szklarczyk et al. 2019; Bleker et al. 2024). Maximum up to three steps of interactions were considered for the primary network construction. BNA was conducted separately for each experimental condition. The input data included gene expression levels, TFs, and associated genes derived from protein-protein interaction (PPI) data. TFs that had binding sites within the expressed genes under the specific experimental conditions were incorporated into the BNA. For any TF-TG interaction model, a TF was included if it had binding sites within the gene promoter regions. This analysis was repeated for each experimental condition, resulting in TF:TG associations for the expressed genes during the major steps of the process.

Given that TFs and their associated PPI components are key factors in TF active binding, a more comprehensive approach was taken using structural equation modeling (SEM). This approach, previously applied successfully in our study investigating miRNA-RBP interactions (Pradhan et al., 2021), allows for the analysis of complex relationships within the regulatory network. SEM consists of two main components: the response (dependent variable) and the independent variables. The primary goal of SEM is to estimate the potential TFs involved at each step. As the data (expression levels of TFs and associated genes from PPI) were continuous, a multivariate gaussian distribution was assumed for all nodes throughout the study. The basic model used was a p-dimensional random vector (X = (X_1_, X_2_, …, X_p_)) with a joint distribution (P= (X_1_, X_2_, …, X_p_)). Here, (X_1_, X_2_, …, X_p_) represent the nodes in the network corresponding to TFs and associated genes from PPI. Bayesian networks (BNs) are directed graphical models where the edges represent conditional independence constraints implied by the joint distribution of X:

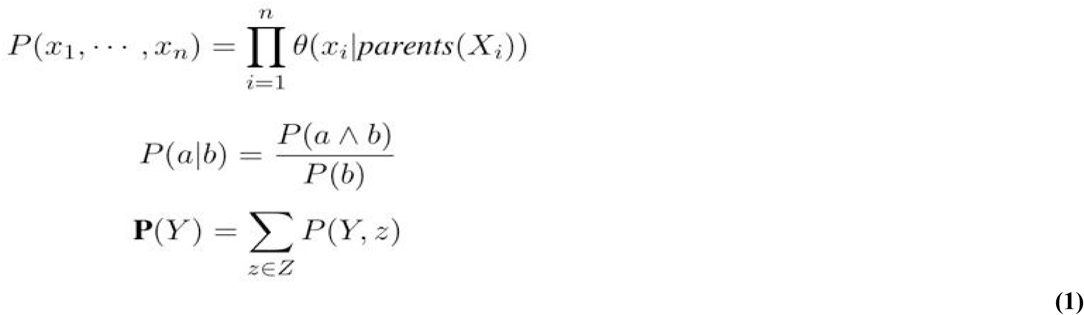

where (x_i_) denotes the parent set of (X_i_) and theta represents the parameters defining the conditional probability distribution (CPD) for (X_i_). The steps for constructing the TF model are as follows with their mathematical implementation:

**1. DAG Estimation:** Directed Acyclic Graphs (DAGs) were estimated for TFs and their PPI partners using gene expression data and TF binding sites as priors, optimized via Hill Climbing (HC) with the Bayesian Information Criterion (BIC) score.
**2. Significant DAG Identification:** Significant DAGs linking TFs to TGs and PPI partners were selected using a statistical test (e.g., chi-square).
**3. SEM Modeling:** Significant DAGs were modeled with a Structural Equation Model (SEM), assuming multivariate Gaussian data, where *X* =*B^T^ X* + *E*, B is the weighted adjacency matrix, and *E N* (0 *, W* ^2^).
**4. Sparse Regularization:** To address high-dimensionality (n<<p), SEM was fitted with a LASSO penalty, controlled by a regularization parameter *λ*.
**5. Parameter Estimation:** Edge weights and regression coefficients were estimated using Maximum Likelihood Estimation (MLE) to fit the observed data.
**6. Conditional Variance Calculation:** The weighted adjacency matrix B = est(β) was used to refine parameter estimates and compute conditional variance.

*Ω* = diag(est(*W*_1_)^2^,…,est(*W_p_*)^2^) was applied as a variance matrix and combining [est(*B*),est(*Ω*)] to calculate the variance covariance matrix Σ and accuracy for each parameter. A suitable convergence criteria (Error tolerance < 10^−4^), precision value > = 85% and an alpha threshold of 0.05 were considered for the selection of the parameters. These parameters decided how larger the effect size (positive/negative) was between TGs and TFs. Detailed mathematical implementations of BNA are provided in **Supplementary File 1**.

The algorithm followed in BNA was performed using “bnlearn”, and “ccdr Algorithm”, both are modules of R (Scutari et al. 2009). The associated basic steps followed in the current approach are described in **Figure 2**.

**Figure 2:**
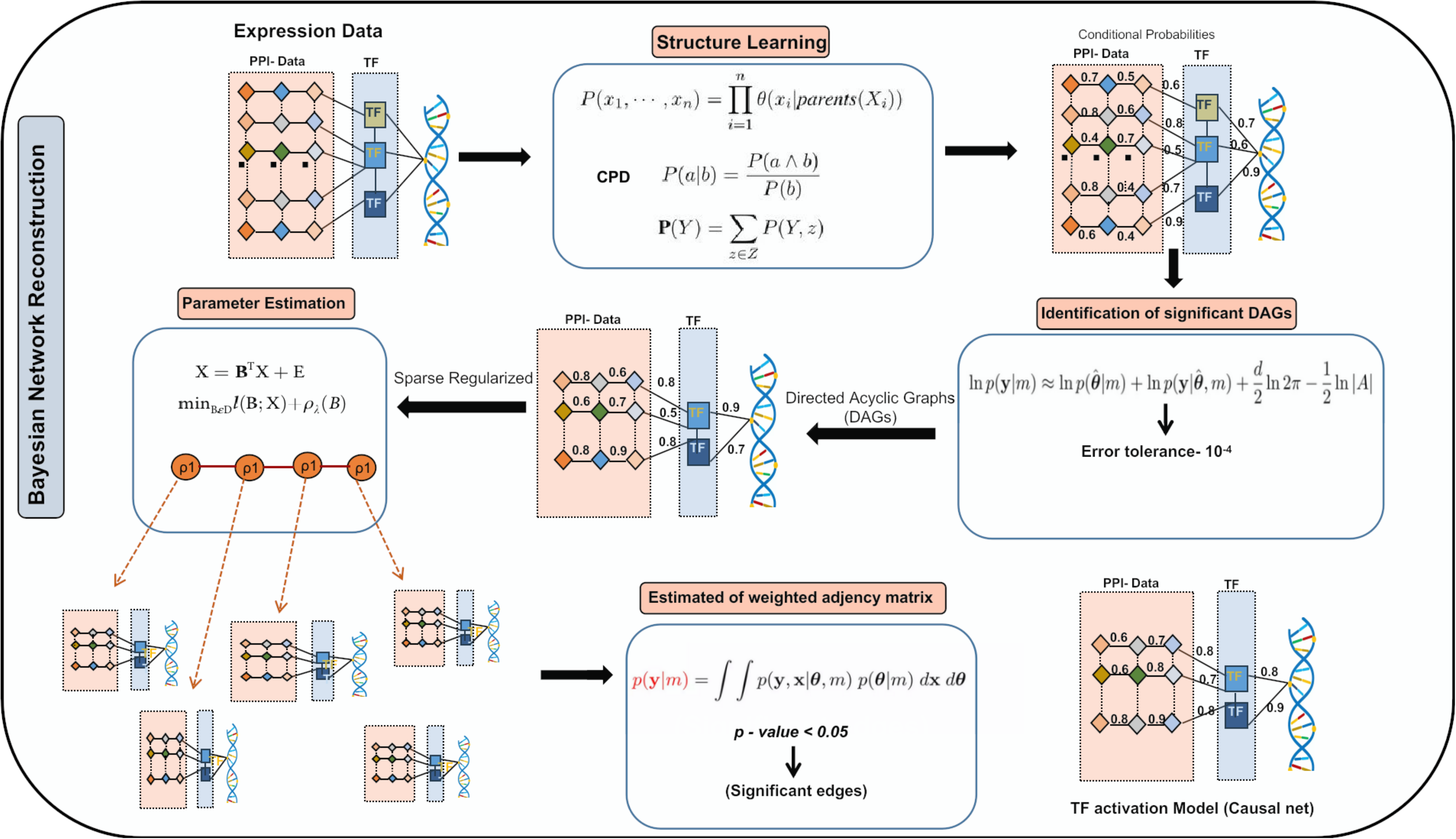
Steps in Bayesian network reconstruction. The process involves: **1)** Estimation of Directed Acyclic Graphs (DAGs) between target genes (TGs), transcription factors (TFs), and their protein-protein interaction (PPI) partners using expression data and prior knowledge. **2)** Identification of significant DAGs based on convergence criteria. **3)** Modeling of significant DAGs with a generalized linear model assuming a multivariate Gaussian distribution. **4)** Parameter estimation between TGs, TFs, and PPI partners using sparse regularization. **5)** Selection of the optimal regularization penalty based on different rho lambda values. **6)** Final parameter estimation using the optimal penalty and criteria-based parameter selection.

### Investigation of Temporal-Varying Networks in Response to Abiotic Stress in *Arabidopsis*

This study deeply explores the biological implications of temporal-varying networks (TVNs) in *Arabidopsis* under abiotic stress conditions, leveraging BNA to uncover dynamic regulatory mechanisms. Abiotic stress, particularly temperature-related stress, has garnered significant attention across scientific, agricultural, and industrial fields due to the increasing frequency of climate and weather extremes driven by global warming. The Intergovernmental Panel on Climate Change (IPCC) Special Report (2024) projects with high confidence that global temperatures will likely rise by 1.5°C between 2030 and 2052 if current warming trends persist, underscoring the urgency to understand and mitigate the impacts of climate extremes.

Using curated dataset, BNA was employed to construct TVNs capturing temporal changes in gene regulatory interactions in response to abiotic stress. To elucidate the functional implications of these networks, gene ontology (GO) (Consortium, 2018) and kyoto encyclopedia of genes and genomes (KEGG) (Kanehisa and Goto, 2000) pathway enrichment analyses were performed using Fisher’s exact test with Benjamini-Hochberg correction (Benjamini and Hochberg, 1995) via the statsmodels library in Python (Seabold and Perktold, 2010). Additionally, BNA-derived TF networks (TF-TG) were compared against the literature-based *Arabidopsis* Transcriptional Regulatory Map (ATRM) (Jin et al., 2016), ATRM encompasses 1,432 TF-TG interactions, to evaluate the concordance between predicted and experimentally validated regulatory relationships. This integrative approach not only highlights key biological functions associated with abiotic stress responses but also assesses the accuracy of the constructed networks by leveraging prior experimental evidence.

### Analyzing TF-cooperativity in regulatory networks

We developed a comprehensive framework for analyzing TF cooperativity using transcriptional regulatory networks constructed from RNA-seq data and promoter sequence analysis. To build the regulatory network, we employed Bayesian network inference to identify relationships between TF-TGs under varying conditions and time points. Promoter regions (2kb upstream) were scanned for TF binding motifs using the PTFspot software tool (Gupta et al., 2024), providing evidence for direct physical interactions between TFs and their target genes.

TF cooperativity was evaluated based on two primary criteria: **1)** expression correlation and **2)** target gene overlap. Expression correlation between TF pairs was quantified using the Pearson correlation coefficient:

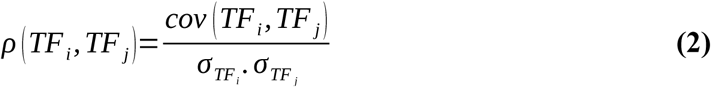

where *cov* (*TF_i_, TF _j_*) denotes covariance, and *σ* represents the standard deviation. A t-test was conducted to assess the significance of the correlation, and p-values were corrected using the Benjamini-Hochberg procedure to control the false discovery rate (FDR). Pairs with a high correlation (*ρ*> 0.7) and a significant (FDR < 0.001) were considered strongly correlated cut off used earlier in similar studies (Zaborowskin et al. 2020; Zhang et al. 2021).

We then measured the overlap between TF target gene sets were measured using the Jaccard Index (J):

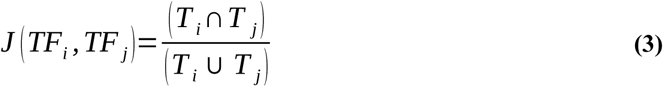

where *T _i_* and *T _j_* denote the sets of genes targeted by *TF_i_* and *TF _j_*, respectively. The statistical significance of the overlap was assessed using the hypergeometric test. Pairs with a high Jaccard index (J > 0.3) and a significant FDR (> 0.01) were identified as significantly overlapping.

### Causal network data for learning causality in deep learning

Condition-specific bayesian causal networks were constructed using the RNA-Seq data and prior known PPI data from the STRING and AGRIS database. The detailed methodology for Bayesian network inference is described in the above sections. Briefly, these networks represent causal relationships between TFs and their interacting proteins, where edges indicate potential regulatory influences or physical interactions. We constructed 11,556 causal networks, covering the 110 TFs across the 46 abiotic stress conditions. These networks, along with associated network statistics (e.g., number of nodes, edges, network density), are made available in our CTF-BIND database (https://hichicob.ihbt.res.in/ctfbind/).

### TFBS identification and dataset creation for TF-DNA interaction module

To construct a high-confidence dataset for the TF-DNA interaction module, we integrated the results from PTFSpot runs with *in vivo* ChIP-Seq data, creating distinct positive and negative datasets for building of our deep learning (DL) model. The 3D structures of 110 TFs were modelled using AlphaFold2, which guided the PTFSpot analysis. Promoter regions, defined as the 2kb upstream of the transcription start site (TSS) for each gene, were extracted to represent core regulatory elements. PTFSpot and PTF-Vac (Gupta et al., 2024) employed these regions with their 3D TF structures to identify potential TFBSs, enabling precise identification of regulatory motifs and enhancing the dataset’s reliability for training the TF-DNA interaction deep-learning model.

#### ChIP-Seq validation (Dataset “A”)

PTFSpot and PTF-Vac detected binding sites were considered a true positive (positive dataset) if they overlapped with a ChIP-Seq peak for the same TF. The sequences corresponding to these overlapping regions were extracted. To capture flanking sequence context, each motif was extended by 75 bases in both the 5’ and 3’ directions, resulting in fixed-length sequences. Because a motif might occur multiple times within a single peak, the number of instances often exceeded the number of peaks. In addition to positive dataset, creating a reliable negative dataset was crucial. Random genomic sequences are not idea ones, as they may not represent realistic non-binding sites. Therefore, we generated negative instances by selecting genomic regions that: **(1)** did not overlap with any ChIP-Seq peaks in any available experimental data, and **(2)** contained motifs similar to those identified by PTFSpot, also with 75bp flanking regions. This approach generated negative examples which were similar to potential binding sites but did not exhibit *in vivo* binding based on ChIP-Seq data. Such negative instances capture the natural scenario of binding preferences where over importance of binding motif is downplayed. The same philosophy has been used previously by the founding algorithm of RBPSpot, (Sharma et al. 2021) a progenitor of PTFSpot/PTF-Vac philosophy. Positive and negative instances were then combined in a 1:1 ratio for each TF. This dataset was used for training and testing the TF-DNA binding DL module. This dataset has been called Dataset “A” throughout this study.

#### Datasets “B” and “C”

Two additional datasets derived from Dataset “A”, “B” and “C”, were created to develop, train, and test the generalized model for TF-DNA interactions. Dataset “B” incorporates gene expression data but disregards the directionality of the edges from the bayesian nets, effectively treating the networks as undirected. On the other hand, Dataset “C” also incorporates gene expression data of the associated network nodes but this time, it considers the directionality of edges in the causal networks. All datasets “A”, “B”, and “C” included data for all the 110 TFs. The ratio of positive to negative data was kept as 1:1 for both the datasets “B” and “C”.

### Word representations of sequence data

The Transformer encoder-decoder architecture is particularly powerful due to its ability to process input and output sequences of varying lengths in a unified, end-to-end manner. This flexibility makes it well-suited for tasks that involve translating or transforming sequences, such as DNA sequences or natural language phrases. Each sequence data were represented using a word-based approach, decomposing each sequence into overlapping k-mers. Specifically, monomer (1-mer), dimer (2-mers), trimer (3-mer), pentamers (5-mers), and heptamers (7-mers) were employed, providing distinct levels of information: monomer captures single base density, dinucleotides capture compositional and base-stacking properties, trimer provide insights into codon adjustment of DNA, pentamers offer insights into local shape, and heptamers reflect regional motif characteristics (Zhou et al., 2013; Sharma et al., 2021; Černý et al., 2020). For example, the sequence ATTGGCAG would be represented by the 2-mers AT, TT, TG, GG, GC, CA, and AG. Given the four-letter DNA alphabet, this approach generates 4 unique 1-mers, 16 unique 2-mers, 64 unique 3-mers, 1024 unique 5-mers, and 16,384 unique 7-mers. Each instance of these k-mers within the input sequences (159-162 bases in length, resulting in a maximum of 469 words) was converted into a unique integer token. These tokens were then transformed into numerical vectors (embeddings) to serve as input to the Transformer encoder. Each k-mer representation (monomer, dimer, trimer, pentamer, and heptamer) was treated independently, with distinct token sets.

### Implementation of the sequence transformers

Tokenized sequences were converted into two-dimensional embedding matrices of size *d × n,* where *d* represents the encoding vector size (32) and *n* the number of tokenized words. Positional information, crucial for sequence understanding, was incorporated by adding positional encodings of dimension *d*, generated using sinusoidal functions (Vaswani et al., 2017), this was combined with the word embeddings, forming the input M’ to the Transformer encoder. Within the multi-headed self-attention mechanism, the embedded sequence M’ was projected onto query (Q), key (K), and value (V) matrices using learnable weight matrices *W^Q^*,*W ^K^*,*W^V^*:

Q=M’. *W ^Q^*, K=M’. *W ^K^*,V=M’. *W^V^*

The attention scores were then computed through scaled dot-product attention between Q and K, followed by softmax normalization and multiplication with V:

This computes pairwise attention weights between all words, capturing their contextual associations.

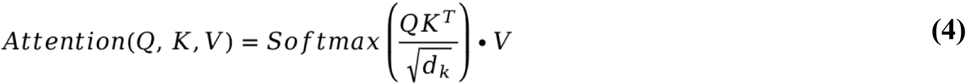

A multi-headed Transformer with 14 attention heads performed this operation in parallel, concatenating the resulting outputs to capture diverse relational perspectives. These combined contextual representations were then processed through a feed-forward network, followed by normalization, dropout regularization, and global average pooling. Model training utilized the Adam optimizer (Kingma and Ba, 2015) with binary cross-entropy loss. This multi-headed self-attention architecture effectively learned long-range dependencies and contextual TF-DNA binding preferences from the tokenized and embedded DNA sequences across 110 *Arabidopsis thaliana* TFs. We went with eight encoder layer. This sequence transformer part output is flattened and concatenated with the Graph transformer output, which was then passed into the second transformer encoder block which derived the hidden features and their relationships which got structured, on which classification could be done in much superior manner. The related detail information about optimization towards the final model is provided in **Supplementary File 1 and Supplementary Table 3 Sheet 1**.

### Graph Representation for the Graph Transformer

The Graph Transformer integrates three components derived from condition-specific causal networks of 110 TFs under 46 abiotic stresses (drought, cold, heat, salinity):

**1. Graph Structure:** Directed acyclic graphs (DAGs) represent regulatory interactions for each TF-condition pair, encoded as adjacency matrices 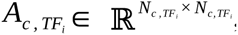, where edges denote causal relationships **(Supplementary File 1: Mathematical Formulation)**.
**2. Node Features:** Nodes (proteins/TFs/target genes) are annotated with condition-specific gene expression, forming feature matrices 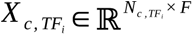, where F represents the z-score normalized gene expression value.
**3. Node-Level Metrics:** Centrality measures (degree, betweenness, eigenvector, closeness, PageRank) computed via NetworkX (Hagberg et al. 2008) quantify structural importance, enriching node representations.

This triad (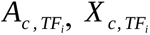, centrality metrics) enables the model to hierarchically learn local-global regulatory patterns while preserving causal semantics. Full computational workflows for adjacency matrix construction and centrality calculations are detailed in **Supplementary File 1.**

### Implementation of Graph-Transformer bimodel

Traditional GNN-based approaches suffer from limitations in aggregating neighboring node vectors, hindering the identification of critical nodes and the inference of long-range structural relationships (Zhang et al. 2022). Graph Transformers are an advanced class of neural networks that extend the transformative self-attention mechanism, originally developed for sequence-based data, to graph-structured data (Yu et al. 2020). Traditional Transformers excel in capturing long-range dependencies within sequential inputs, and Graph Transformers adapt this capability to graphs by enabling nodes to attend to other nodes across the network. This architecture effectively captures both local and global relationships, making it highly suitable for analyzing graph-based structures such as TF causal networks.

### Graph Transformer Architecture for Condition-Specific TFBS Identification

The Graph Transformer models TF causal networks, where nodes represent TFs or target genes, and edges denote regulatory interactions. This approach captures both biological properties and network topology to enhance the identification of TF binding sites (TFBS).

#### Node and Edge Embedding Initialization

Each node in the TF causal network is initialized with a feature vector integrating gene expression data and centrality metrics (degree, betweenness, and closeness centrality) to reflect its functional and structural importance. Similarly, edge embeddings encode the nature of regulatory interactions (e.g., activation or repression) using a learnable mapping function, allowing the model to differentiate between distinct regulatory roles.

#### Attention mechanism

The Graph Transformer employs a scaled dot-product attention mechanism to model the relationships between nodes and their neighbors in the TF causal network. The attention score between a pair of nodes **“u”** and **“v”** is computed as:

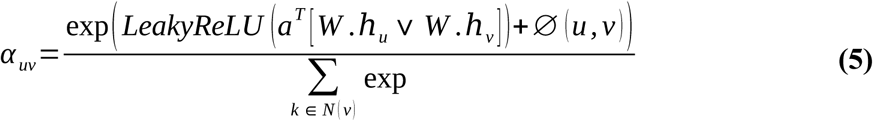

where, W is a learnable weight matrix used to transform node embeddings, **a** is a learnable vector used for scoring attention,[.∨.]denotes concatenation of two vectors,∅(*u, v*) represents the spatial encoding (e.g, shortest path distance or a predefined measure of relatedness between nodes u and v),*N* (*v*) denotes the set of neighbors of node v.

#### Node representation update

The node embedding at layer *l* +1is updated as:

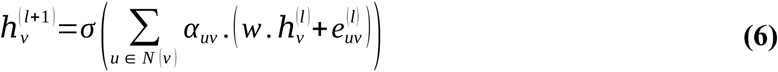

where: *σ* is a nonlinear activation function (e.g., ReLU), *α_uv_* is the attention coefficient, 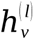 is the node embedding of u at layer l, 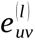 is the edge embedding at layer l.

#### Edge representation update

The edge embedding *e_uv_* is updated at each layer to encode dynamic interactions as:

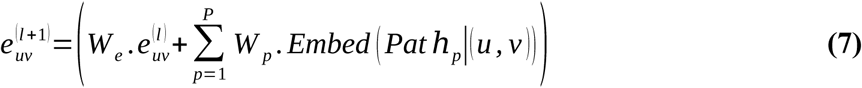

where: *W_e_* is a learnable transformation matrix for edge embeddings,*Pat ℎ_p_* (*u, v*) encodes the *p-th* shortest path between nodes u and v, *Embed*(.) maps the path into a vector representation, *W _p_* is a learnable weight matrix for the *p-th* path.

We implemented the Graph Transformer using PyTorch Geometric, employing 12 Transformer layers to process 160-dimensional node embeddings. Each layer incorporated multi-head attention (8 heads, 20-dimensional projections per head), leveraging shortest-path encoding (N = 3) to capture both local and global regulatory patterns. Skip connections and layer normalization mitigated over-smoothing, ensuring stable learning. Node embeddings were dynamically refined using nonlinear transformations (ReLU) and attention-weighted aggregation, while edge embeddings incorporated multi-hop interactions to model regulatory influence. The output was flattened, concatenated with sequence-based features, and passed through a secondary Transformer block to integrate cross-modal relationships. For classification, the model utilized a feed-forward network with dropout (0.2), LeakyReLU activations, and global average pooling, followed by a single-node sigmoid layer. The Adam optimizer (learning rate: 0.015) was used with binary cross-entropy loss, optimizing over 45 epochs (batch size: 32).

The transformer network extracts and structures hidden features and their relationships, making classification more effective. Since the present problem in this study was not translation but classification, decoders were not needed and instead the encoder output were taken as input for next step of extreme gradient boosting. For this purpose the output of the transformer was passed to the XGBoost classification part. XGBoost was the choice as it has consistently scored highest in Kaggle benchmarking studies and performs exceptionally good on structured and curated features sets. **Figure 3** provides a snapshot of how this entire system is working. Additional architectural details, including edge update equations and optimization, are available in **Supplementary File 1.**

**Figure 3:**
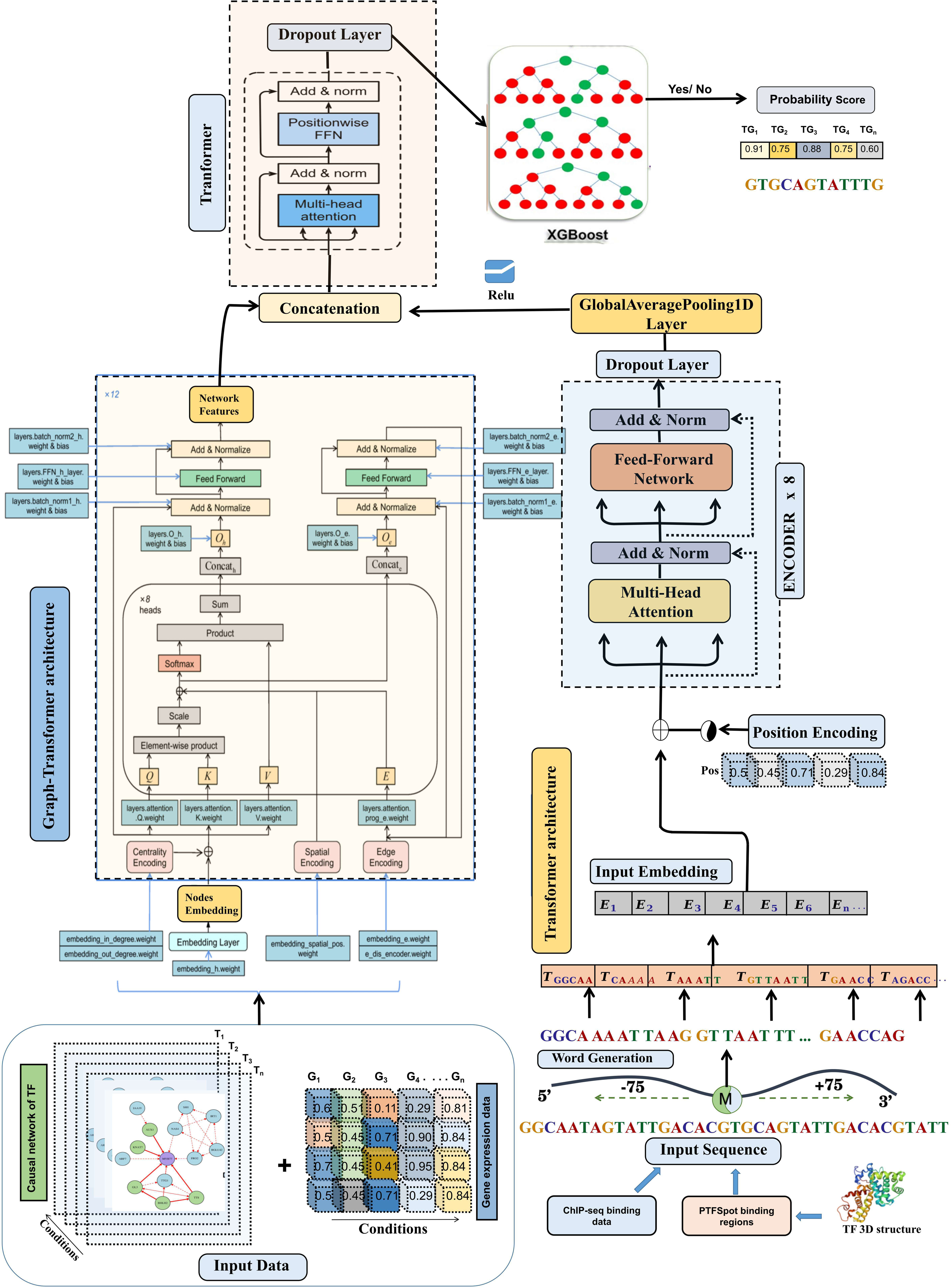
Overview of the architecture and workflow of the CTF-BIND framework for predicting transcription factor (TF) binding activity. Input data includes TF causal networks, gene expression profiles, ChIP-seq data, TF 3D structures, and promoter sequences. A Graph-Transformer processes network features, while a Transformer encodes DNA sequence information. Features are concatenated, pooled, and passed through an XGBoost classifier to predict TF binding probabilities for target genes (TGs) under specific conditions. This system enables accurate TF binding predictions directly from transcriptome data, eliminating the need for additional ChIP-seq experiments.

### Performance Evaluation

The Graph Transformer-based model’s performance was rigorously assessed using a comprehensive set of metrics derived from the confusion matrix, true positives (TP), false negatives (FN), false positives (FP), and true negatives (TN), including sensitivity (TPR), specificity (TNR), precision, F1-score, and Matthews Correlation Coefficient (MCC). Sensitivity measures the model’s ability to correctly identify positive instances, while specificity evaluates its accuracy with negative instances. Precision indicates the proportion of true positives among all predicted positives, and the F1-score, a harmonic mean of precision and recall, provides a balanced metric for imbalanced datasets. MCC, a indicator accounting for all confusion matrix elements, was used to evaluate overall consistency, with higher scores reflecting strong performance across classes, especially in imbalanced scenarios. Additional metrics, such as the Area Under the Receiver Operating Characteristic Curve (AUC/ROC) and mean absolute error (MAE), were computed to assess the sensitivity-specificity trade-off and average detection error, respectively. To ensure reliable and unbiased evaluation, datasets were split into 70% training and 30% testing sets, with models built on the training portion and evaluated on the unseen testing portion. This process was validated through 10-fold randomized trials, where the dataset was repeatedly divided into 70:30 splits, ensuring no overlap between training and testing instances across folds. This rigorous approach prevented memorization and over-fitting, maintaining the integrity and consistency of the performance assessment.

Performance measures were done using the following equations:

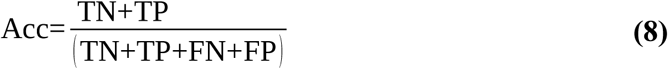

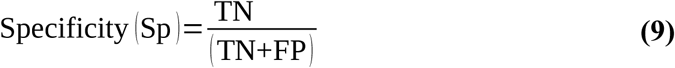

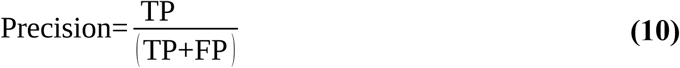

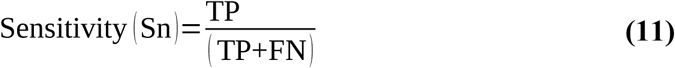

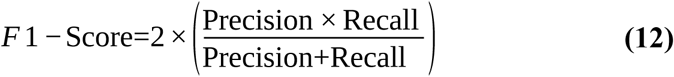

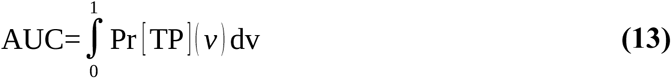

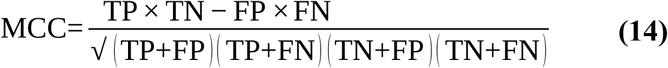

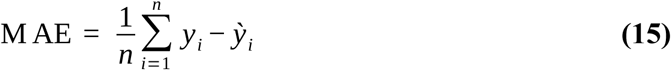

Where:

TP = True Positives, TN = True Negatives, FP = False Positives, FN = False Negatives, Acc = Accuracy, AUC = Area Under Curve. MAE = Mean Absolute Error

## Results and discussion

### TF binding information forms the root Causal Nets of spatio-temporal regulation

A total of 490 ChIP-seq samples for 124 TFs were obtained from the ChIP-Hub, PlantPan 3.0, and GEO databases, encompassing approximately 9.8 TB of reads, representing 17 experiments. These reads underwent rigorous quality checks and filtering, followed by analyses to identify target genes. Additional target genes were identified using PTFspot, which utilized the 3D structure of all TFs along with a 2 kb upstream region of each gene’s promoter. The identified TF-target gene (TF-TG) interactions from these approaches overlapped by approximately 80%, resulting in a final set of the most confident TF-TG interactions for downstream analyses. The distribution of binding sites for each TF is illustrated in **Supplementary Figure S1**, with detailed information provided in **Supplementary Tables S6**. Notably, the number of binding sites for TFs varied significantly, influenced by the total number of experimental conditions reported. Some TFs were studied multiple times, contributing to a higher count of identified binding sites.

The bubble chart (**Supplementary Figure S1**) displays the diverse binding patterns of TFs to target genes, with each bubble representing a TF and its size corresponding to the number of genes bound. DREB2B and RAP2.9 emerged as the TFs with the highest number of gene interactions, binding to 783 and 465 genes, respectively. These TFs are critical for abiotic stress responses, particularly in regulating genes involved in drought and salinity tolerance (GO:0009651, FDR < 0.001) (Liu et al., 2020; Zhao et al., 2016). STZ (Salt Tolerance Zinc finger protein) also exhibited significant representation, binding to 690 gene targets. This aligns with STZ’s known role in osmotic stress regulation and its involvement in high salt condition responses (Zhang et al., 2018). TFs such as MYB83, WRKY8, and ERF38 displayed substantial numbers of bound genes, with 564, 514, and 254 targets, respectively. The MYB family is well-documented for its roles in abiotic stress tolerance, particularly in secondary metabolite production and reactive oxygen species (ROS) scavenging pathways (GO:0000302, FDR < 0.0046) (Dubos et al., 2020). Similarly, WRKY and ERF families are essential for modulating stress responses through hormone signaling pathways, including abscisic acid (ABA) signaling (Fujimoto et al., 2000; Liu et al., 2024). A smaller cluster of TFs, including HSFA2, bHLH071, and IAA29, exhibited relatively fewer gene interactions, with binding sites ranging from 44 to 148 genes. These TFs have more specific roles in stress signaling (GO:0140467, FDR < 0.001), targeting a narrower set of genes that are tightly regulated under stress conditions (Zhao et al., 2016). Several TFs, such as TCP3, PRR5, and NF-YC2, exhibited a broader spectrum of target genes across various conditions, indicating their involvement in diverse biological processes and stress adaptations. For instance, TCP3 is known for its role in cell proliferation and growth, while NF-YC2 is part of the NF-Y family, which mediates stress responses by regulating key stress-responsive genes (Hao et al., 2019). The analysis reveals significant variability in the number of genes bound by each TF, with certain TFs exhibiting exceptionally high numbers of targets due to their central roles in abiotic stress pathways. TF such as DREB2B, STZ, and RAP2.9 demonstrated significant involvement in stress-responsive gene networks (GO:0006950, FDR < 0.0016), highlighting their potential as key regulators. Notably, among these, only DREB2B has been previously characterized as a master regulator in abiotic stress responses (Zuo et al., 2020). After this finding STZ and RAP2.9 also need to be evaluated.

This analysis established a crucial foundation for identifying the most promising TF:TGs interaction pairs. The subsequent application of Bayesian Network Analysis (BNA) further provided the reasoning circuits for these interactions, selectively filtering for associations with functional relevance. This refinement was based on high-confidence evidences, integrating both gene expression and PPI interaction network data, ensuring the most biologically meaningful TF:TGs associations to evolve and come up.

### Bayesian networks derive functional TF-TG associations with their Causal routes capable to explain the spatio-temporal regulation

ChIP-seq data served as the foundational dataset to identify TF-TG associations, providing binding evidence for TFs and their target genes (TGs). The functional binding of TFs have impact on the expression of the downstream gene in a spatio-temporal manner To reveal their functional associations and uncover causal relationships, Bayesian Network Analysis (BNA) was applied. TF interactions and their influence on target genes are often context-dependent, involving dynamic regulatory networks. The causal network construction began with the primary TF-TG associations derived from ChIP-seq data, supplemented with target gene identification from PTFspot, which considers structural information of TFs with the sequence context properties of the target regions.

Protein-protein interaction (PPI) data for the identified TFs were incorporated to establish their primary static networks. These static networks were then enhanced with condition-specific dynamic information by integrating RNA-seq expression data for network components, TFs and their target genes across various experimental conditions. The BNA methodology, detailed in the methods section, was employed to derive directionality and causality, ensuring that the identified associations were biologically meaningful and functionally relevant.

Approximately 96% (6,180) of the TF-TG combinations identified through BNA demonstrated strong correlations (|r| ≥ 0.8) with a statistically significant p-value of 0.007, as determined by a binomial test, thereby supporting the reliability of the inferred causal networks. A total of 6,438 TF-TG interactions were identified, spanning 874 unique transcription factors and 3,561 target genes across all experimental conditions. These networks captured key regulatory pathways for abiotic stress responses. For example, TFs such as DREB2A, NAC019, MYB60 and WRKY8, previously identified as master regulators **(Supplementary Figure S2-3)**, were shown to orchestrate large networks of stress-responsive genes.

The identified causal networks also highlighted condition-specific regulatory dynamics. For instance, in drought conditions, DREB2A exhibited strong associations with genes involved in osmotic adjustment and dehydration tolerance (GO:0009414, FDR < 0.001), while NAC019 displayed causal regulation of oxidative stress-responsive genes (GO:0006979, FDR < 0.001). Similarly, during salinity stress, STZ (Salt Tolerance Zinc Finger protein) and WRKY8 emerged as central regulators, modulating genes involved in ion homeostasis and ABA signaling (GO:0009738, FDR < 0.001). More than 91% of the TF-TG combinations identified through BNA demonstrated consistent positive or negative associations when validated under individual experimental conditions.

### Causal analysis of transcription factor networks in response to Abiotic stress in *A. thaliana*

This study investigates the causal regulatory pathways within time-varying TF networks in *A. thaliana* under cold, heat, drought and salt stress, utilizing BNA to reconstruct and analyze dynamic TF-target gene (TG) relationships. BNA uses time-series gene expression data to detect causal activation patterns, revealing how TFs are sequentially activated and subsequently bind to target genes to propagate stress responses across the network.

Time-series gene expression data from seven time points (T0–T6) for cold stress and five time points (T0–T4) for heat stress and seven time points for drought and five time points for salt were analyzed. Differentially expressed genes (DEGs) were identified at each time point relative to the initial state (T0). A total of 47, 33, 642, 2,662, 6,119, and 7,166 DEGs were detected for cold stress from T1-T6. A total of 1,093, 515, 1,595, and 7,243 DEGs were found for heat stress from T1-T4. A total of 997, 425, 1,616, 1,945, 3,435 and 7,243 DEGs detected for drought stress from T1-T6. A total of 93, 215, 1,935, and 943 DEGs detected for salt stress from T1-T4. BNA constructed causal networks for each time point, capturing TF-TG interactions that drive stress responses.

The analysis revealed modular structures in the networks, where clusters of genes, activated in a sequential manner, demonstrated causal relationships. Modules activated at earlier time points triggered the activation of neighboring modules at later stages, indicating a cascading regulatory effect. Cold stress networks exhibited a delayed response, with significant activation beginning at T3, while heat stress networks showed immediate responses starting at T1. similarly drought and salt stress networks showed immediate responses starting at T1. This delay in cold stress responses concerns well with previous studies on AtGenExpress datasets (Wanke et al., 2010a).

Prominent TFs known to regulate stress responses were identified in the networks. For cold stress, CBF3, a key regulator initiating cold-responsive gene expression (Medina et al., 2011), appeared at T3 **(Supplementary Figure S3)**. TFs shared between cold and drought stress, such as DREB2A and DREB2B, were observed in T3 networks, reflecting overlapping regulatory mechanisms (Xiong et al., 2002). Heat shock factors (HSFs) like HSFA2 and HSFC1, which regulate heat-shock-related proteins, were also found in T3 and T4 cold stress networks, highlighting their roles in mitigating protein misfolding and maintaining cellular homeostasis during both conditions (Swindell et al., 2007). In heat stress networks, well-documented TFs such as HSFA2, known for regulating downstream heat-responsive genes (Schramm et al., 2006) and ERF family TFs (Mizoi et al., 2012) were prominently featured **(Supplementary Figure S3)**.

For drought stress, early response regulators such as bZIP TFs (e.g., bZIP1 and bZIP5), known for their roles in ABA-mediated signaling and antioxidant system regulation (Liu et al. 2023; Zhu et al. 2018), were observed at T1, reflecting their involvement in rapid stress adaptation (e.g., osmotic adjustments and oxidative stress mitigation). Similarly, AP2/ERF TFs and MYB TFs, including ODORANT1, appeared in early networks (T2-T3) **(Supplementary Figure S4)**, orchestrating the expression of stress-tolerance genes linked to water retention and root development. NAC TFs, such as NAC022, were notable in T4 networks, enhancing ABA-dependent pathways to regulate ion homeostasis and water balance under prolonged drought stress (Hong et al. 2016). For salinity stress, late-response TFs such as WRKY TFs and zinc finger proteins featured prominently in T2-T3 networks, reflecting their role in maintaining long-term stress tolerance through transcriptional modulation and post-translational gene silencing. Additionally, RAV TFs (e.g., RAV1 and RAV2) were detected in T4 networks **(Figure 4)**, for water-loss mitigation and seed germination regulation (Ghorbani et al. 2019). Heat shock factors like HSF TFs and C2H2 zinc finger TFs were also observed in salinity stress networks, highlighting their roles in protecting cellular homeostasis and inducing heat shock proteins during sustained stress conditions.

**Figure 4:**
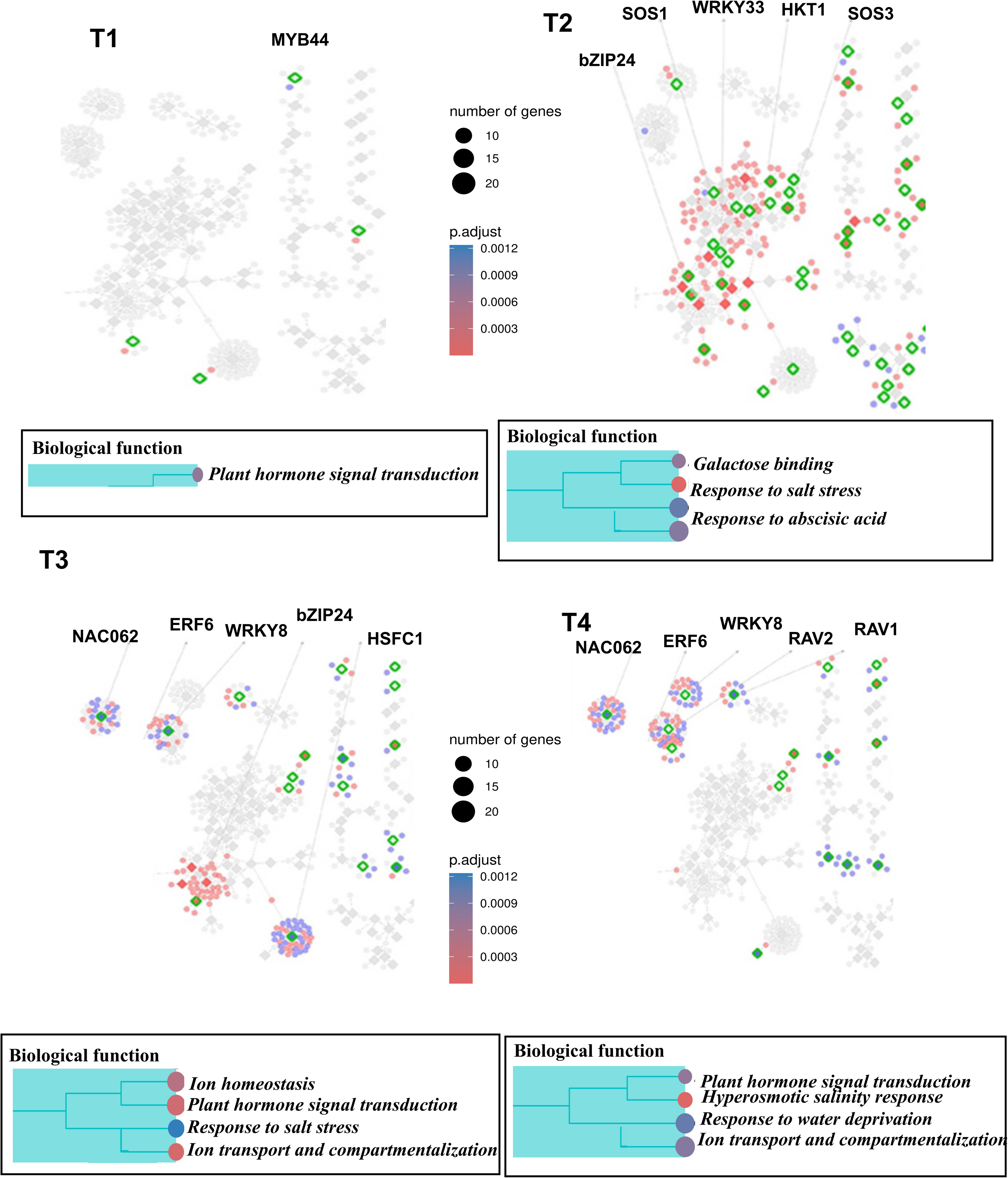
Dynamic Time-varying network of different time points (T1 ∼ T6) for Salinity stress experiment data in *Arabidopsis*. Red and blue nodes represent up/down DEGs, and green border rhombuses represent the identified regulatory TFs. Tables show top-2 enriched terms for each of KEGG pathways [K] and GO terms [G] with adjusted p-values by Benjamini-Hochberg correction. Some interesting TFs are named in the figure.

Gene Ontology (GO) and KEGG pathway enrichment analyses further highlighted the biological relevance of these networks. Cold stress networks were enriched for terms like “response to cold” and “response to water” and the “plant hormone signal transduction” pathway, consistent with known cold stress mechanisms **(Supplementary Figure S2)** (Eremina et al., 2016). Heat stress networks were enriched for the GO term “response to chitin” and the KEGG pathway “hormone signal transduction,” both linked to heat stress responses in prior studies (Eremina et al., 2016) **(Supplementary Figure S3)**. Drought stress networks were enriched for GO terms such as “response to water deprivation” and “regulation of stomatal movement,” reflecting mechanisms critical for maintaining water balance under osmotic stress and KEGG pathway “plant hormone signal transduction” was significantly enriched **(Supplementary Figure S4)**. Salinity stress networks were enriched for GO terms like “response to salt stress” and “ion homeostasis,” which are essential for mitigating ionic toxicity and osmotic imbalances and KEGG pathway “ion transport and compartmentalization” emerged as a key pathway **(Figure 4)**, emphasizing the importance of regulating ion uptake and intracellular ion concentration during salinity stress.

To validate the causal TF-TG relationships, BNA networks were compared to the literature-based *Arabidopsis* Transcriptional Regulatory Map (ATRM) (Jin et al., 2016). Among the 380 edges in the cold stress network and 478 in the heat stress network, 543 in the drought stress network and 432 in the salt stress network 92,76,81 and 67 edges, respectively, were supported by prior experimental evidence. Significant overlap with ATRM (p < 0.00147 for cold stress; p < 0.0098 for heat stress; p < 0.00561 for drought stress; p < 0.00123 for salt stress) underscores the reliability of BNA based identifications made here. Specific cold stress edges, such as DREB2A → LTI78 (Xiong et al., 2001) and CBF3 → ERD10 (Seki et al., 2002), and heat stress edges, such as HSFA9 → HSP101 (Kotak et al., 2007) and WRKY39 → MBF1C (Li et al., 2010), and drought stress edges, such as bZIP5 → ERF3 (Liu et al. 2023) and NAC022 → ERF144 (Wang et al., 2022), and salt stress edges, such as RAV1 → HSP71 (Ghorbani et al. 2019) and HSFA2 → WRKY70 (Zhao et al., 2017), were among those supported by literature. These findings demonstrate the ability of BNA to elucidate causal regulatory mechanisms in abiotics stress, highlighting key TFs and pathways that orchestrate the plant’s adaptive response through dynamic TF-TG interactions.

The insights gained from causal TF networks provide a foundation for developing stress-resilient crops. By identifying key TFs like DREB2A, bZIP1, and HSFA2 that orchestrate early and sustained responses to multiple stresses, targeted genetic engineering or marker-assisted breeding can enhance plant resilience. For instance, overexpression of drought-responsive TFs such as NAC022 could improve water retention and reduce drought-induced yield losses. Similarly, modulation of salt stress regulators like RAV1 may enhance salinity tolerance in crops grown in coastal or saline-prone regions.

In addition to enhancing individual stress tolerance, understanding shared regulatory pathways can facilitate the development of multi-stress-resistant crops. For example, simultaneous manipulation of common TFs like DREB2A could provide tolerance to cold, drought, and salt stress. Furthermore, the enriched GO and KEGG terms such as “response to water deprivation,” “ion homeostasis,” and “plant hormone signal transduction” highlight critical pathways that could be targeted to optimize stress response efficiency while maintaining plant growth and productivity. In this study, the exact chains of composing genes, and their directionality have been revealed.

### Causal Regulatory Paths in Abiotic Stress Response

Plants are constantly exposed to a variety of abiotic stresses, such as heat, cold, drought, and salinity, which challenge their survival and productivity. To overcome these challenges, plants have evolved intricate regulatory networks that orchestrate a coordinated response, ensuring resilience and continued growth in fluctuating environmental conditions.

To unravel the complex causal interactions behind plant responses to these stresses, we applied Bayesian network analysis. This method revealed causal relationships between genes, TFs, and other regulatory components in the system, providing insight into the pathways that govern plant stress tolerance. Through this analysis, dynamic stress-specific and common regulatory modules were identified, furthering our understanding of plant resilience mechanisms.

### Stress-Specific Sub-Networks

#### Cold Stress

The cold stress network highlighted the central role of the CBF regulon in cold acclimation, with CBF1, CBF2, and CBF3 TFs acting as key regulators. Additionally, CBF-independent mechanisms involving RAV1, ZEP, and MYBL2 were found to contribute to cold tolerance **(Figure 5-i)**. The integration of circadian and light signaling pathways, mediated by LHY1 and CCA1, further modulated cold stress responses. This indicates that cold stress responses are governed by a highly integrated network of TFs, balancing both immediate and long-term adaptive responses.

**Figure 5:**
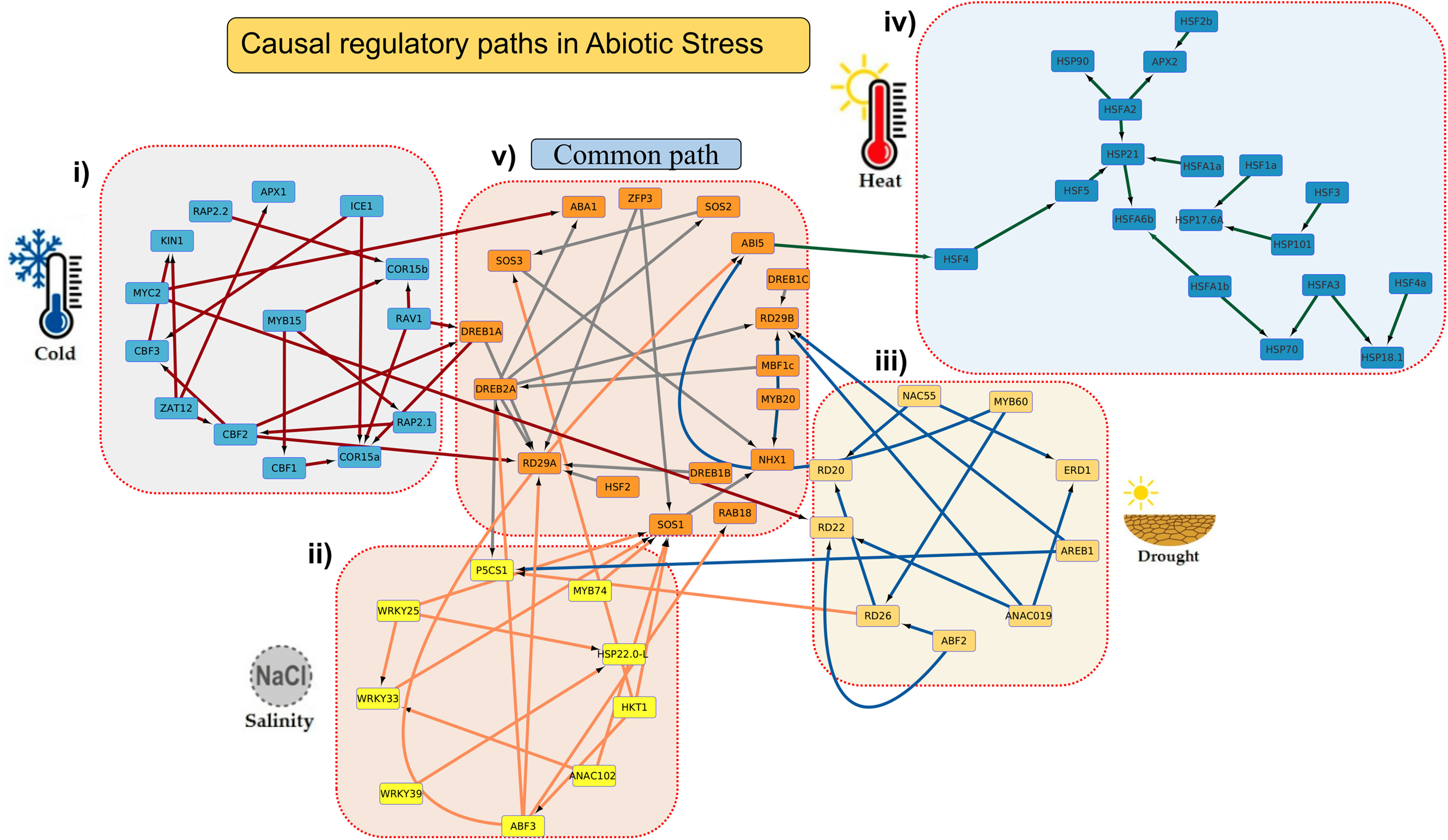
Causal regulatory pathways of transcription factors (TFs) under abiotic stress conditions. **(i)** Cold stress network: Key TFs such as ICE1, MYC2, and ZAT12 regulate downstream targets, forming an interconnected network with nodes like RAP2.2 and CBFs driving cold-specific responses. **(ii)** Salinity stress network: Salinity-responsive TFs (e.g., WRKY25, NAC102, and ABF3) interact through pathways involving ion transporters (e.g., SOS1) and osmotic regulation genes (e.g., P5CS1). **(iii)** Drought stress network: Drought-regulated TFs, including NAC55, MYB60, and ABF2, mediate pathways controlling dehydration-responsive genes such as RD22 and AREB1. **(iv)** Heat stress network: Heat-shock factors (e.g., HSFA2, HSFA6b) coordinate responses involving heat-shock proteins (e.g., HSP70, HSP101) to maintain protein stability under heat stress. **(v)** Common pathways: Shared nodes like DREB2A and ABA1 connect multiple stress responses, illustrating the integration of signaling pathways across different abiotic stresses. This figure highlights the condition-specific and shared regulatory mechanisms underlying plant responses to abiotic stress.

#### Heat Stress

In the heat stress sub-network, HSFA2 was identified as a key regulator of the heat shock response (HSR), activating heat shock proteins (HSPs) to protect cellular proteins under high temperatures. Negative regulators such as HSFB1 and HSFB2b helped fine-tune this response **(Figure 5-iv)**, while the DREB2 TF family mediated the expression of heat and drought-responsive genes. The multifunctional coactivator MBF1c was also crucial in linking TFs to transcriptional machinery, ensuring an efficient response to heat stress.

#### Salt Stress

The salt stress sub-network revealed TIFY10A as a critical regulator, with JAZ5 acting as a co-repressor in the salinity stress response. PHYTOCLOCK 1 (PCL1) was found to play a significant role in modulating circadian rhythms, helping plants optimize their responses to salt stress. Additionally, WRKY TFs (WRKY33 and SOS2) were central to regulating ion transport and detoxification, further complicating the salt stress response **(Figure 5-ii)**.

#### Drought Stress

The drought stress network identified DREB2A as a central player in regulating osmotic adjustment and water conservation. MYB60 was critical for stomatal regulation, while WRKY22 integrated drought and salinity responses **(Figure 5-iii)**. The involvement of PIF3 in ROS homeostasis further highlighted the multifaceted nature of drought tolerance mechanisms.

#### Common Regulatory Pathways Across Stresses

This analysis revealed a “common path” connecting all abiotic stress conditions, indicating a shared regulatory architecture. Central nodes such as DREB2A, HSP90, and RD29A were frequently observed, suggesting their universal roles in abiotic stress tolerance **(Figure 5-v)**. Specifically, We found that SOS2, ZFP9, and other ABA-responsive elements orchestrated responses to osmotic stress in both drought and salinity, confirming ABA’s central role as a hormonal regulator. The shared presence of HSP90 in heat and drought stress pathways indicated its importance in stabilizing cellular proteins and preventing denaturation under various stress conditions **(Figure 5-v)**. Regulators like PIF3 and ZAT12 modulated ROS homeostasis, contributing to stress adaptation across different abiotic challenges.

This Bayesian network analysis provided valuable insights into the complex and dynamic regulatory landscape governing plant responses to abiotic stress. The identified stress-specific and common regulatory modules demonstrated the diverse yet interconnected strategies plants employ to cope with environmental challenges. The frequent occurrence of key regulatory interactions such as DREB2A, HSP90, and RD29A, as visualized in the network **(Figure 5-v),** emphasizes their essential roles in achieving stress tolerance. These findings not only deepen our understanding of plant stress biology but also offer potential targets for genetic engineering and breeding strategies aimed at improving crop resilience in the face of increasingly challenging environmental conditions.

### Condition-Specific TF Cooperativity in Abiotic Stress Responses

Our analysis of TF cooperativity within transcriptional regulatory networks revealed distinct and biologically significant patterns of cooperative interactions under different abiotic stress conditions. By integrating expression correlation, target gene overlap, and functional enrichment analyses, we identified key cooperative TF pairs mediating stress responses. A total of 43 significantly cooperative TF pairs (correlation coefficient ≥ 0.7, Jaccard index ≥ 0.3, FDR < 0.001) were identified, with stress-specific distributions: 14 pairs in heat, 8 in drought, 11 in salt, and 10 in cold stress conditions.

#### Cold Stress

Under cold stress, we observed strong cooperativity between CBF1 (C-repeat Binding Factor 1) and ICE1 (Inducer of CBF Expression 1). These TFs exhibited a high Pearson correlation coefficient (ρ = 0.92, FDR < 0.001) and a significant Jaccard index (J = 0.45, FDR < 0.01) **(Figure 6)**, indicating both strong expression correlation and substantial target gene overlap. GO enrichment analysis of the overlapping target genes revealed enrichment of terms related to “response to cold” (GO:0009409, FDR = 1.2e-5) and “cold acclimation” (GO:0009628, FDR = 8.7e-4). This aligns with previous studies demonstrating the synergistic action of CBF1 and ICE1 in regulating cold-responsive genes (Chinnusamy et al., 2003). ICE1 activates CBF expression, which then bind to CRT/DRE elements in cold-responsive gene promoters. Our analysis successfully captured this known cooperative relationship.

**Figure 6:**
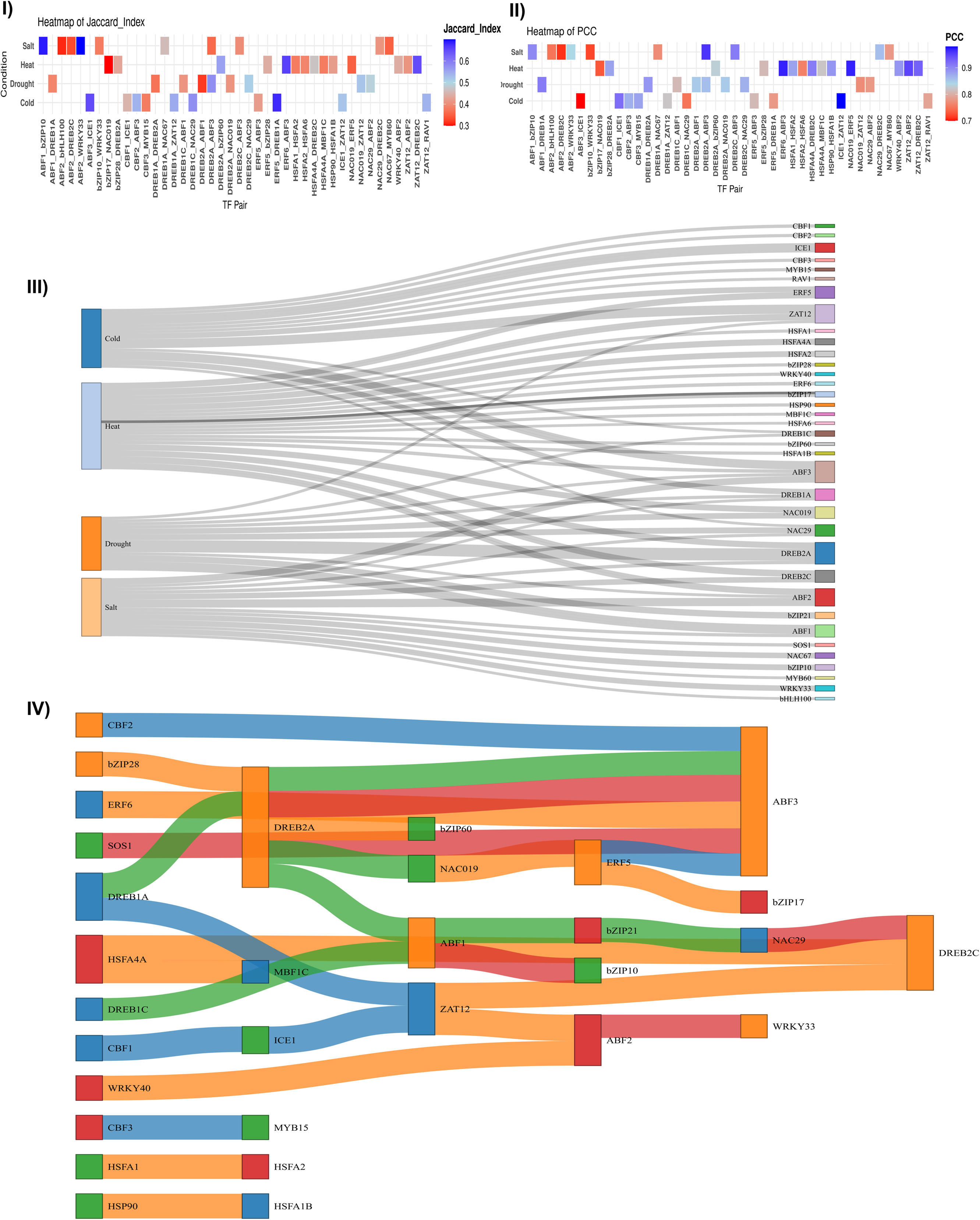
Transcription Factor (TF) cooperativity in response to various abiotic stresses: **I)** Heatmap depicting the Jaccard Index, which represents the overlap of target genes between TF pairs. Higher Jaccard Index values indicate a greater overlap in target genes. **II)** Heatmap illustrating the Pearson Correlation between TF gene expression and their target genes across different stress conditions. **III)** Sankey diagram showing the distribution of TFs across abiotic stresses, including Heat, Cold, Drought, and Salinity. Each TF is linked to specific stress categories, indicating its involvement in multiple stress responses. **IV)** Sankey plot illustrating cooperative TF pairs targeting shared genes. Each colored strip represents a different stress type (Heat, Cold, Drought, Salinity), demonstrating how TF pairs collaborate under various abiotic conditions.

#### Heat Stress

In response to heat stress, we identified strong cooperativity between HSFA1 (Heat Shock Transcription Factor A1) and HSFA2 (Heat Shock Transcription Factor A2), validating previously reported interactions (Liu et al., 2021). These TFs showed a high expression correlation (ρ = 0.88, FDR < 0.001) and a significant target gene overlap (J = 0.38, FDR < 0.01) **(Figure 6)**. The HSFA1-HSFA2 cooperative pair showed sequential activation, with HSFA1 peaking at 1 hour and HSFA2 at 3 hours post-stress. Their shared target genes (n=143) were significantly enriched in processes including “protein folding” (GO:0006457, FDR = 1.2e-12) and “response to heat” (GO:0009408, FDR = 3.4e-9). This is consistent with the established role of HSFA1 and HSFA2 in regulating HSP expression, crucial for protein folding and preventing aggregation under heat stress (Scharf et al., 2012). A novel cooperative interaction was also identified between HSFA4A and MBF1C (correlation = 0.82, Jaccard index = 0.38) **(Figure 6)**, with 87 shared target genes enriched in “cellular response to oxidative stress” (GO:0034599, FDR = 2.1e-7), suggesting a link between heat stress and oxidative stress responses.

#### Drought Stress

Under drought stress, a cooperative interaction between was identified DREB2A and NAC019 (correlation = 0.85, Jaccard index = 0.41), demonstrating strong cooperation **(Figure 6)**. This pair showed coordinated regulation of 112 genes involved in “response to water deprivation” (GO:0009414, FDR = 4.3e-8) and “ABA signaling” (GO:0009738, FDR = 1.7e-6), consistent with their roles in ABA-dependent drought tolerance (Yoshida et al., 2010). The previously identified cooperativity between DREB2A and ABF1 was also confirmed in our analysis (ρ = 0.85, FDR < 0.001 and J = 0.32, FDR < 0.01), reinforcing their role in osmotic stress responses through regulation of genes like RD29A and LEA1.

#### Salinity Stress

Under salinity stress, cooperative interaction between was identified NAC67 and MYB60 (correlation = 0.78, Jaccard index = 0.35) **(Figure 6)**, suggesting a functional association. This pair showed coordinated regulation of 85 genes involved in “response to salt stress” (GO:0009651, FDR = 2.1e-6) and “response to osmotic stress” (GO:0006970, FDR = 8.5e-5). This aligns with the known roles of NAC and MYB transcription factors in abiotic stress responses, including salinity tolerance (Kumar et al., 2017; Hoang et al., 2020). NAC TFs often regulate downstream genes involved in stress responses, while MYB TFs can modulate various processes, including secondary metabolism and stress responses. The observed co-expression and shared target genes suggest a potential synergistic effect of NAC67 and MYB60 in mediating salinity tolerance. Further analysis identified potential novel TF cooperativity pairs under salinity stress. One such pair is bZIP10 and WRKY33 (correlation = 0.72, Jaccard index = 0.38) **(Figure 6)**. These TFs coordinately regulate 32 genes enriched in “response to oxidative stress” (GO:0006979, FDR = 5.7e-4) and “cellular response to chemical stimulus” (GO:0070887, FDR = 1.2e-3). Salinity stress often induces oxidative stress, and the coordinated action of bZIP and WRKY TFs, known for their roles in stress signaling and defense responses, could be crucial for mitigating the damaging effects of salt. bZIP TFs are often involved in ABA signaling and stress responses, while WRKY TFs play a crucial role in plant immunity and abiotic stress responses (Banerjeeet al., 2002; Liu et al., 2024). Their co-regulation of genes related to oxidative stress suggests a coordinated defense mechanism under saline conditions. Another potential cooperative pair identified was AREB1/ABF2 and bHLH100 (correlation = 0.75, Jaccard index = 0.31) **(Figure 6)**. This pair showed coordinated regulation of 58 genes involved in “abscisic acid-activated signaling pathway” (GO:0009737, FDR = 3.9e-5) and “response to abscisic acid” (GO:0009738, FDR = 9.2e-6). ABA plays a crucial role in mediating plant responses to various abiotic stresses, including salinity. AREB/ABF TFs are key regulators of ABA-dependent gene expression, while bHLH TFs are involved in various developmental and stress-related processes (Dong et al., 2024; Lei al., 2011). The observed co-regulation of ABA-responsive genes suggests a potential synergistic effect of AREB1/ABF2 and bHLH100 in modulating ABA signaling under salinity stress.

These results highlight the condition-specific and shared nature of TF cooperativity in response to abiotic stresses. Our analysis identified known cooperative TF pairs and uncovered novel interactions, providing functional insights into their roles in stress responses. The integration of expression correlation, target gene overlap analysis, combined with GO enrichment and motif analysis, provided a robust framework for identifying and characterizing TF cooperativity. The observed synergistic actions of these TFs underscore the importance of combinatorial regulation in fine-tuning gene expression under stress conditions. The temporal shifts in TF network structure and gene expression underscore the dynamic nature of stress responses. The identification of stress-specific interactions suggests both dedicated and general stress response mechanisms.

### Value of Bayesian Network Analysis (BNA) in revealing causal regulatory dynamics

While RNA-seq and co-expression networks identify correlated gene expression patterns, Bayesian Network Analysis (BNA) adds critical value by inferring directional causal relationships and conditional dependencies within transcriptional networks. Unlike co-expression networks, which merely highlight associations, BNA models how perturbations (e.g., abiotic stress) propagate through regulatory pathways, distinguishing direct drivers from indirect effects. For instance, in cold stress, BNA revealed the causal path ICE1 → CBF1 → cold-responsive genes, clarifying ICE1’s upstream regulatory role in activating CBF1-a relationship obscured in co-expression networks that show only correlation (ρ = 0.92). Similarly, BNA uncovered sequential activation in heat stress (HSFA1 peaking before HSFA2), highlighting temporal hierarchies invisible to static co-expression analyses.

BNA also exposed novel, condition-specific cooperativity missed by traditional approaches. For example, the interaction between HSFA4A and MBF1C under heat stress, linked to oxidative stress responses, was identified through causal paths involving shared target genes and upstream regulators. Co-expression alone could not disentangle whether this interaction was causal or coincidental. Similarly, BNA resolved the DREB2A-NAC019 partnership in drought stress as a driver of ABA signaling, distinguishing it from spurious correlations.

By integrating expression data with prior knowledge (e.g., TF binding motifs, GO terms), BNA provided mechanistic insights into how cooperative TF pairs regulate stress responses. For instance, the bZIP10-WRKY33 interaction under salinity stress was validated through causal links to oxidative stress genes, explaining their synergistic role in mitigating salt-induced damage.

### Causal networks are capable to profess accurately about the condition specific TF-binding, when learned “deeply”

Causal networks have demonstrated a remarkable capacity to accurately identify condition-specific transcription factor binding when trained with deep learning methodologies. These networks, as previously outlined, serve as frameworks for reasoning about the intricate regulatory circuits that govern stress-induced, context-dependent TF binding events. By integrating experimentally validated binding data, sequence-specific information, and condition-specific transcriptome profiles, these causal networks enable the deciphering of spatio-temporal regulatory dynamics. This identify power suggests that such networks could be leveraged to infer system-wide conditions solely from transcriptome data, offering a transformative approach to understanding gene regulation.

Building on this foundation, the study advanced to the next phase: training these causal networks to enhance their predictive accuracy and explanatory potential. This process involved synthesizing multiple layers of biological information, including TF binding data validated through experimental assays, DNA sequence motifs, and transcriptomic profiles reflective of specific physiological states. The methodology employed in this integration, along with its detailed implementation, is comprehensively described in the Methods section and further elaborated in the associated supplementary materials **(Supplementary File 1)**. This systematic approach laid the groundwork for developing a computational system capable of modeling complex regulatory interactions with unprecedented precision.

Central to this advancement is the development of the PTFSpot/PTFVac software system, a highly reliable tool designed to pinpoint TF binding sites across the genome with exceptional accuracy. This system integrates three critical components: 3D structure of the TF, the most probable binding motif or “seed” as determined by the RBPSpot algorithm (Sharma et al. 2021), and the contextual influence of flanking genomic regions. While PTFSpot/PTFVac excels at identifying potential TF binding sites, determining the precise conditions under which binding occurs at these sites traditionally necessitates experimental techniques such as Chromatin Immunoprecipitation sequencing (ChIP-seq). However, conducting ChIP-seq across the vast array of possible biological conditions is impractical, and until recently, no computational alternative has adequately addressed this challenge.

To overcome this limitation, a novel Graph-Transformer-based deep learning framework was employed to unify these disparate factors—TF structure, binding motifs, flanking contexts, and condition-specific expression data—into a cohesive DL model. This approach culminated in the creation of a pioneering computational tool, the first of its kind, capable of identifying condition-specific TF binding events with high efficiency and specificity across all potential genomic binding sites. By harnessing the power of causal networks and deep learning, this model not only predicts where a TF is likely to bind but also elucidates when such binding is biologically relevant, based solely on computational analysis. A detailed exposition of this methodology, including the software’s architecture and performance metrics, is provided in a dedicated **Supplementary File 1.**

#### Context sequence information form the solid foundation

Accurate identification of TF binding sites necessitates considering the sequence context surrounding the core binding motif. As detailed in the Methods section, both the *in silico* PTFspot results and the *in vivo* ChIP-seq data provide such contextual information alongside the primary binding motif. The identification of these core motifs is crucial for selecting relevant contextual features, as demonstrated in our previous work. While these motifs are significantly enriched in binding regions, they are not exclusive to them and can also occur in non-binding regions of the genome. Therefore, these discovered motifs served as anchor points for identifying potentially relevant interaction sites across the DNA, emphasizing the importance of analyzing their surrounding sequence context. To capture this context, we included 75 bp of flanking sequence on both the 5’ and 3’ ends of each motif. This approach is supported by previous studies demonstrating the effectiveness of flanking regions in capturing local environmental information relevant to nucleic acid interactions. This context can encompass various factors, including the presence of co-occurring motifs, sequence- and position-specific biases, and structural/shape characteristics of the DNA. These contextual features serve as powerful discriminators between true binding sites (positive instances) and non-binding regions (negative instances). By incorporating these flanking regions, we constructed the datasets used in this study (Datasets “A,” “B,” and “C”), as detailed in the Methods section.

To understand the impact of various features in the TF:DNA interaction model, an ablation study was conducted. It evaluated 15 different combination of word representation strategies for the building of model. This analysis aimed to assess the contribution of each representation, in conjunction with TF-associated network information using a Graph Transformer, to distinguishing between preferred (positive instance) and non-preferred (negative instance) binding regions. Dataset “A” (positive instances derived from ChIP-Seq data) was partitioned into 70% training and 30% testing sets for this evaluation. Initial results were derived using monomeric, dimeric, and trimeric word representations where words were DNA sequences substrings of single, di and tri mers achieved an accuracy of 72.24%, 73.67%, and 74.94%, respectively on Dataset “A” **(Supplementary Figure S5)**. Subsequent incorporation of pentameric and heptameric representations yielded improved performance, reaching 78.03% and 81.54% accuracy, respectively **(Supplementary Figure S5)**. These representations corresponded to sequence window sizes of 156 and 154 units (pentamers and heptamers). Consequently, we created combined datasets incorporating all five representations (monomers, dimers, trimers, pentamers, and heptamers). This multi-representation approach enabled the Transformer models to learn contextual information with greater information sharing across the representations. This combined sequence representation strategy was then integrated with TF-specific network information within the Graph Transformer framework. The pairing of pentameric and hexameric representations resulted in a significant leap in the accuracy to 87.21%. Following this, the best performance was observed on combining representations of dimeric, pentameric, and hexameric, resulting in an impressive accuracy of 92.51% with a total of 469 representative words. These results emphasize the necessity of contextual information sharing among different representations, as they collectively harnessed their unique attributes to raise a superior model.

As depicted in **Supplementary Figure S5**, the combined representations consistently outperformed any single representation. Detailed information regarding the individual representations and their performance metrics is provided in **Supplementary Table 3 Sheet 1-5**. Implementation details of the optimized Transformer architecture are described in the Methods section and illustrated in **Figure 3**.

#### Transcriptome profile further fills the nodes with additional information and bolster the association graph

The Graph Transformer bimodal demonstrated to capture both local and distal sequence dependencies with Bayesian nets, effectively integrating information across multiple learned feature representations. We randomly selected 10 TF, namely WRKY40, ANAC032, ABI5, ABF2, ZAT10, HAT22, DREB2A, HSFA1A, MYB44, and SEP3 for evaluation on Dataset “A,” the model achieved accuracy 81.7%, indicating low performance across the majority of analyzed TFs **(Figure 7a)**. This highlights the model’s ability to learn discriminatory patterns directly from the sequence context of experimentally validated binding sites. Remarkably, when trained and tested on 10 TFs from Dataset “B,” which incorporates normalized gene expression data along with undirected TF networks, the Graph Transformer bimodal performance further improved to 84.59% accuracy **(Figure 7b)**. This substantial improvement underscores the critical role of gene expression data in enhancing TF binding site detection. Gene expression data provides valuable functional context by reflecting the transcriptional activity of genes within the network. This information allows the model to prioritize binding sites that are more likely to be functionally relevant under specific conditions. By integrating gene expression information, the model can discriminate between potential binding sites that are merely present in the DNA sequence and those that are actively utilized in a given cellular context. This integration effectively filters out spurious binding sites and focuses on those with likely functional consequences, leading to the observed increase in model’s accuracy.

**Figure 7:**
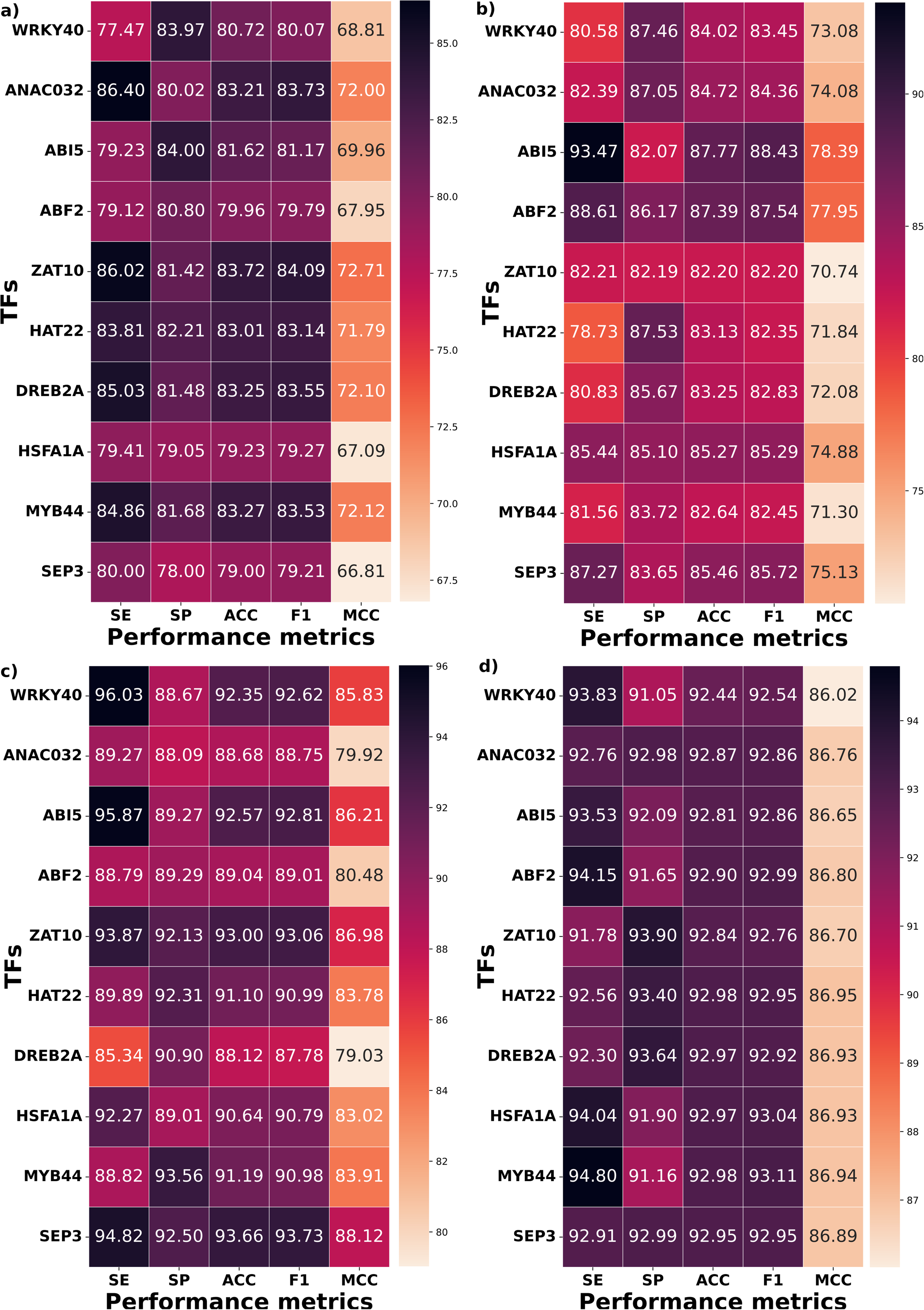
Performance evaluation of transcription factor (TF) binding models across various datasets and configurations. **(a)** Evaluation on Dataset A: Performance assessment using ChIP-seq data, showcasing baseline accuracy and specificity based solely on experimental TF binding profiles. **(b)** Evaluation on Dataset B: Assessment incorporating gene expression data without considering the directionality of regulatory networks**, (c)** Evaluation on Dataset C: Performance evaluation using gene expression data combined with the directionality of causal networks, illustrating significant accuracy gains by leveraging network-based causality. **(d)** Performance of GraphTransformer with XGBoost: Analysis of the GraphTransformer architecture integrated with XGBoost, using Dataset C. This configuration highlights enhanced high capabilities through the combination of deep learning and ensemble methods.

#### BNA derived causality and directed graph boost the capabilities of the model to detect conditional binding by TFs

Furthermore, training and testing on same ten TFs from Dataset “C,” which incorporates normalized gene expression data along with directed causal edges of condition-specific TF networks, resulted in a significant further performance boost, achieving 91.035% accuracy **(Figure 7c)**. This substantial improvement demonstrates the critical importance of incorporating network causality and edge directionality. The directed edges in these networks represent inferred causal relationships between TFs and their target genes, providing crucial information about the flow of regulatory information. The directed edges in causal networks encode interaction weights that quantify both the strength and directionality of regulatory influences, enabling the Graph Transformer to model how perturbations in upstream TF activity propagate through the network to modulate gene expression and binding events. For example, under drought stress, increased activity of DREB2A (upstream) may amplify its regulatory effect on NAC019 (downstream), with edge weights reflecting the magnitude of this influence. By explicitly learning these causal relationships, the model distinguishes active regulatory pathways (e.g., ABA-dependent drought responses) from inactive ones, prioritizing binding events driven by condition-specific signals (e.g., heat-induced HSFA1A activation). The integration of sequence data, transcriptome profiles, and causal networks creates a synergistic framework that filters out non-functional binding sites (e.g., motifs near inactive genes) while highlighting those with biological relevance (e.g., ABF2 binding near ABA-responsive genes). This multi-modal strategy elevates accuracy by aligning identify with the dynamic regulatory logic of living systems, offering unprecedented resolution for studying context-dependent gene regulation in development, stress adaptation.

However, the deep learning system delivered a good accuracy of 91.035% but there was a gap of 0.92% between sensitivity and specificity, though not a big gap, yet we tried to reduce it further. In doing so, the output layer of the transformer having the LeakyReLu activation function was replaced by XGBoost for the classification purpose. XGBoost was the choice as it has come consistently at the top along with deep learning approaches in Kaggle benchmarkings and performs exceptionally good on structured data and manually extracted features input sets. In our case, the Transformer’s encoder became the feature feeder to XGBoost. This hybrid deep-shallow model further reduced the performance gap between the sensitivity and specificity to just 0.79% while also increased the accuracy slightly to ∼93% (**Figure 7d**).

10-fold random trials for the assessment of robustness and consistency was performed concurred with the above observed performance level and scored in the same range consistently. All of ten randomly selected TFs achieved high quality ROC curves with high AUC values in the range of 0.9089 to 0.9407 (Dataset “C”) while maintaining reasonable balance between specificity and sensitivity **(Figure 8a-j)**. Also, the MAE for training was found to be 0.2851, while for testing it was 0.2854, resulting in a very small difference of only 0.003. A t-test comparing the MAE values for the training and test sets yielded a highly insignificant result of ∼38%, much above the significance threshold of 5% or lower, further confirming the absence of any possibility of any significant over-fitting (**Supplementary Table 3 Sheet 5)**. To conduct an unbiased performance testing without any potential recollection of data instances, it was ensured that no overlap and redundancy existed across the data.

**Figure 8:**
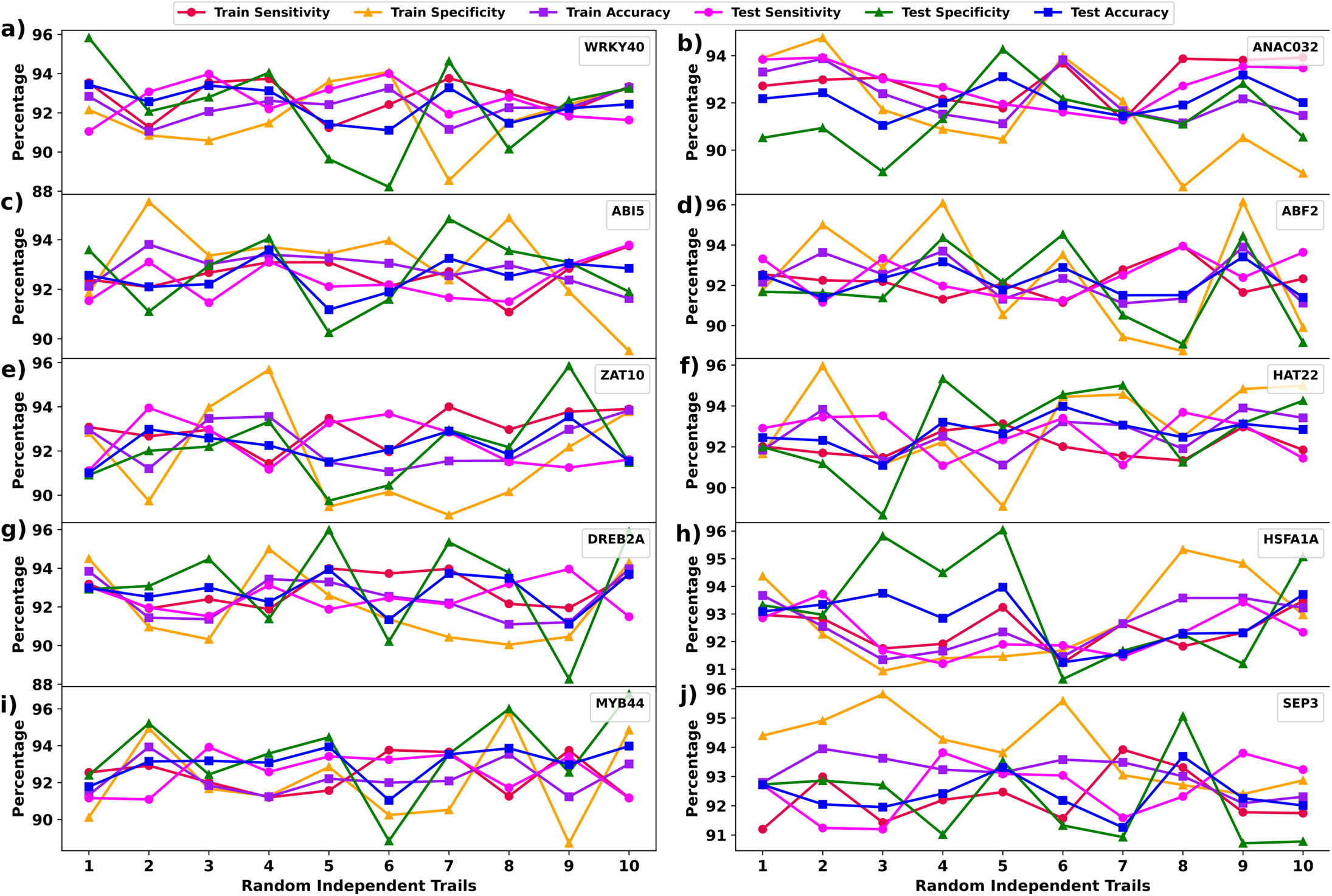
Performance evaluation on randomly selected 10 transcription factors (TFs): The model was tested on 10 randomly selected TFs to assess performance consistency. The results demonstrated a strong balance between specificity and sensitivity, validating the reliability and robustness of the model for TF binding under diverse conditions.

Afterwards, we deployed this DL model system was deployed to the rest 100 TFs from dataset “C” to train and test the model. Each of these TFs were never included in the training of the above mentioned co-learning model. Here, the DL model achieved an astonishing average accuracy of 92.51% with balanced sensitivity and specificity values of 92.84% and 92.18%, respectively (**Supplementary Figure S7**). The remarkable performance consistency ensured about the robustness of the raised transformer models and reliability of its all results. It was evident that the Transformer effectively grasped both distant and nearby words associations, while acquiring knowledge through multiple hidden features. The full architectural details of the optimized Transformer-XGBoost system is illustrated in **Figure 3**.

### Grad-CAM as an Explainable AI Framework for TFBS Identification Using Causal Networks

Understanding TFBS and their regulatory mechanisms is crucial for uncovering gene regulation processes. As deep learning models become increasingly sophisticated in detecting TFBS, the need for explainability grows. To address this challenge, we implemented Gradient Weighted Class Activation Mapping (Grad-CAM). This adaptation of Grad-CAM aims to identify the contextual and gene features in TF causal networks that most significantly contribute to TF binding site discovery under specific conditions. By incorporating causal network information and sequence-level features, this approach enhances the interpretability of complex models and provided insights into the mechanisms governing TF binding. To compute the importance of each node in the TF causal network for identifying DNA binding, we first calculated the gradient of the output y (the likelihood of TF binding) with respect to the node-specific activations. This is expressed as 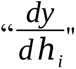 represents the feature vector of node *v_i_* in the network. The importance weight *α_i_* for each node*v_i_* is derived by averaging the gradient components over all dimensions *d* of *ℎ_i_*:

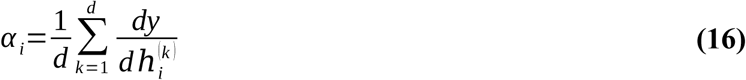

where 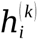 is the *k*-th dimension of *ℎ_i_*. To compute the contribution of node *v_i_*, we used the ReLU activation function was used to retain only the positive contributions, ensuring that the negative gradients did not detract from the importance measure. The node score is given by:

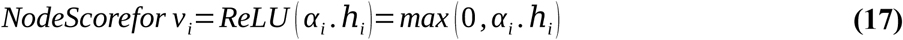

The total contribution of the network was then calculated by summing the ReLU-activated scores across all the nodes in the graph:

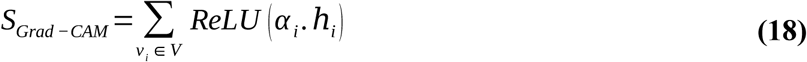

This ensures the distribution of importance across all nodes reflects their relative roles in determining the TF binding under specific conditions.

In our implementation of Grad-CAM, we selected the normalization layer following the XGboost layer as our layer of interest **(Figure 3)**. This layer contains distribution maps for diverse nodes present in the causal network. Adhering to the weighting methodology proposed by Grad-CAM, we quantified the significance of these nodes and computed a weighted summation of all distribution maps. This aggregated map effectively highlights the nodes that are pivotal to binding activities. Subsequently, we conducted enrichment analysis to determine whether each unique k-mer was enriched among all k-mers with the highest average Grad-CAM scores. This analysis was tested across 35 TFs utilized in the study. For user given sequence, the model first computes a Grad-CAM score profile highlighting the contextual features most strongly associated with TF binding status. The relevance of sequence features, such as k-mers, was evaluated by identifying those with the highest average Grad-CAM scores. In the causal network, Grad-CAM was applied to identify key nodes and edges contributing to the performance of the model. The network represents TFs and genes as nodes, with directed edges encoding causal relationships and regulatory influences. By analyzing the Grad-CAM score distributions across the network, nodes with significant contributions to the binding sites were identified.

As an example to show the use of Grad-CAM score to detect various important factors in any given condition binding of TF, we discuss here the case of WRKY33 binding. To identify key regulatory nodes within the TFs causal networks and understand their influence on TF binding, we applied Grad-CAM, visualizing the importance of each node (TF or gene) for the discovery of TF binding to specific DNA sequences under particular conditions. Applying Grad-CAM to the WRKY33 causal network pinpointed key sequence features influencing its binding, with top-scoring k-mers exhibiting 84.94% concordance with known WRKY33 motifs. Five key nodes (WRKY25, MPK3, STZ, MKK6, and MYB51) exhibited high Grad-CAM scores (>0.8), **(Figure 9)** and their *in silico* removal from the network abolished accurate WRKY33 motif identification, demonstrating their crucial role in determining WRKY33 binding specificity. To validate these computational findings, we conducted a literature review and found strong experimental support for the identified interactions: WRKY33 is established as a condition-dependent master regulator in plant specialized metabolism, particularly in the camalexin and 4OH-ICN pathways, coordinating with MYB51 in *Arabidopsis thaliana* for gene expression and metabolic fluxes (Barco et al. 2020; Chen et al. 2024). Both WRKY33 and WRKY25, members of the WRKY family involved in stress responses, exhibit distinct regulatory roles in *Gossypium hirsutum*, with WRKY33 acting as a stronger negative stress regulator (Ehsan et al. 2023; Li et al. 2011). While direct evidence for WRKY33 interactions with MPK3, STZ, and MKK6 was limited, but their known roles in stress response pathways (MPK3 in MAP kinase signaling, STZ in salt tolerance, and MKK6 also in MAP kinase cascades interacting with WRKY TFs) provide compelling contextual support (Andreasson et al. 2005; Mao et al. 2011; Zhou et al. 2020; Han et al. 2019). This integrated approach of computational analysis and experimental validation highlights the complex regulatory mechanisms governing TF binding, demonstrating that WRKY33’s binding is influenced not only by its core motif but also by interactions within its regulatory network, which very ably explain the condition specific binding. The implemented Grad-CAM helps to identify the important factors in providing the mechanistic of TF binding and gene regulation. With this all, the present deep leaning model, CTF-BIND emerges as a first of its kind highly valuable tool to detect condition specific TF binding and a promising alternative to ChIP-seq. It also gives opportunity to discover the casual agents in condition specific TF regulation. This approach have extended to other species and beyond stress specific conditions also.

**Figure 9:**
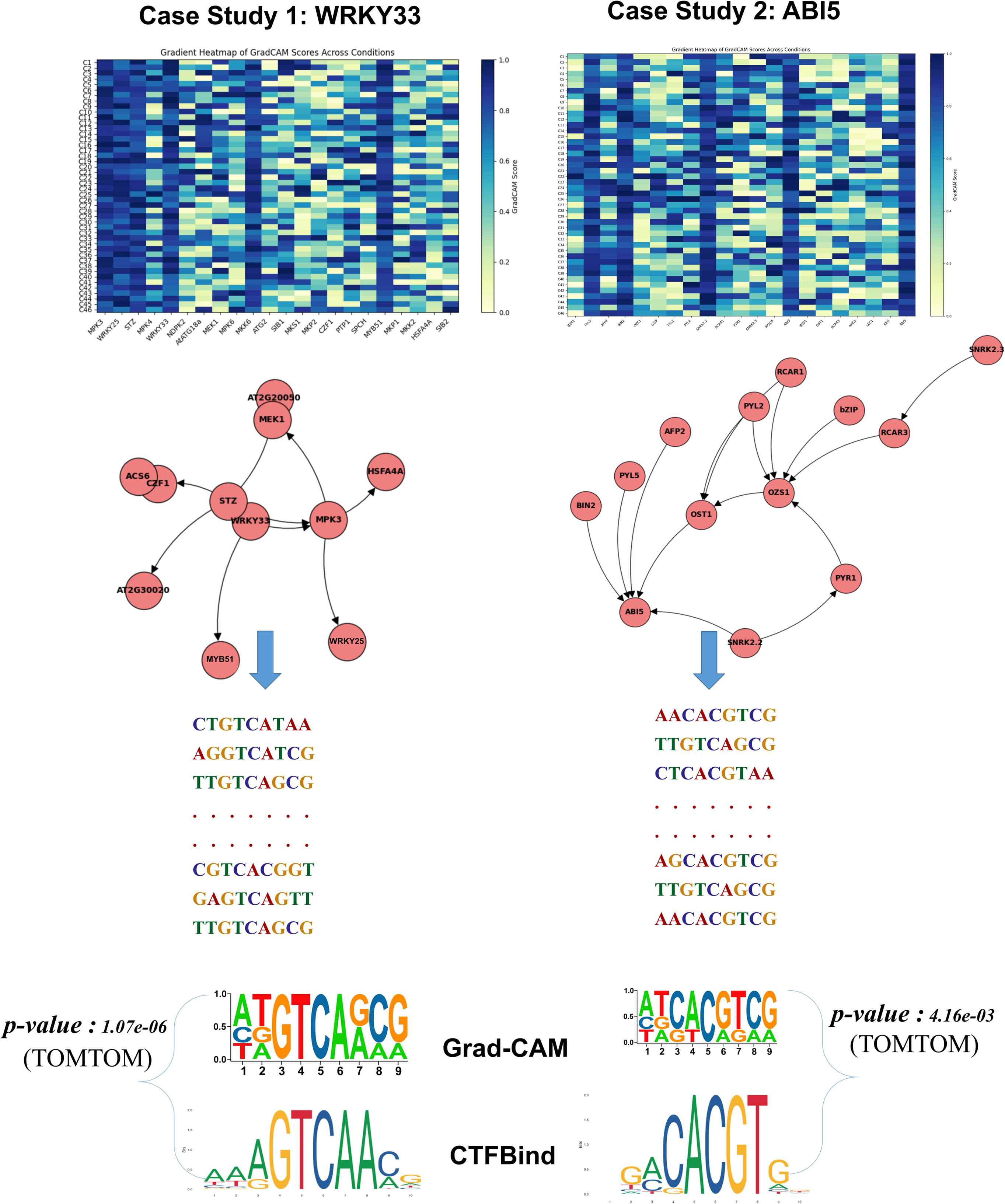
This figure illustrates the application of Grad-CAM to visualize the importance of different features (nodes) within the network for predicting the binding of two transcription factors (TFs): WRKY22 and ABI5. The gradient heatmap highlights the most important features, providing insights into the underlying decision-making process of the model.

### Applying CTF-BIND to Abiotic stress data: A demonstration of conditionality-specific TF binding

As an application of demonstration CTF-BIND tool (https://scbb.ihbt.res.in/CTFBIND/), we investigated condition-specific TF binding to the target gene RD29A (Responsive to Dehydration 29A) in *A. thaliana* under abiotic stress. RD29A, known for its responsiveness to drought, cold, and salt, with varying expression across tissues (Msanne et al., 2011), was selected as the target gene. Using publicly available cold and heat stress gene expression datasets from GEO, we performed differential expression analysis, confirming RD29A’s differential expression across these conditions. The 2kb upstream promoter region of RD29A was extracted, and potential TF binding regions (TFBRs) were identified using PTFSpot **(Figure 10)**. This analysis revealed a set of candidate TFs, including ABF2, that could potentially bind to the RD29A promoter. To determine condition-specific binding, we applied CTF-BIND. Providing the RD29A promoter sequence and gene expression data under cold stress, CTF-BIND discovered ABF2 binding to specific sites within the promoter. Subsequent correlation analysis between RD29A and ABF2 expression levels revealed a strong positive correlation (R = 0.92) in cold stress conditions, suggesting ABF2’s regulatory role in RD29A expression under cold stress **(Figure 10)**. Additionally, RD29A gene expression showed a significant correlation with hub TFs like ABI5 and DREB1A, gene expression within the ABF2 TF network suggesting a broader regulatory network. In contrast, when CTF-BIND was applied using heat stress expression data, it did not identify any ABF2 binding to the RD29A promoter. Consistent with this identification, the correlation between ABF2 and RD29A expression was significantly weaker (R = 0.28) under heat stress **(Figure 10)**. These results demonstrate CTF-BIND’s ability to accurately identify condition-specific TF binding. The observation of ABF2 binding to RD29A under cold stress, but not heat stress, and the corresponding changes in correlation between TF and TG expression provides strong evidence for the tool’s capacity to identify causal regulatory relationships in response to environmental cues as well as capabilities to be an alternative to costly experiments like ChIP-seq.

**Figure 10:**
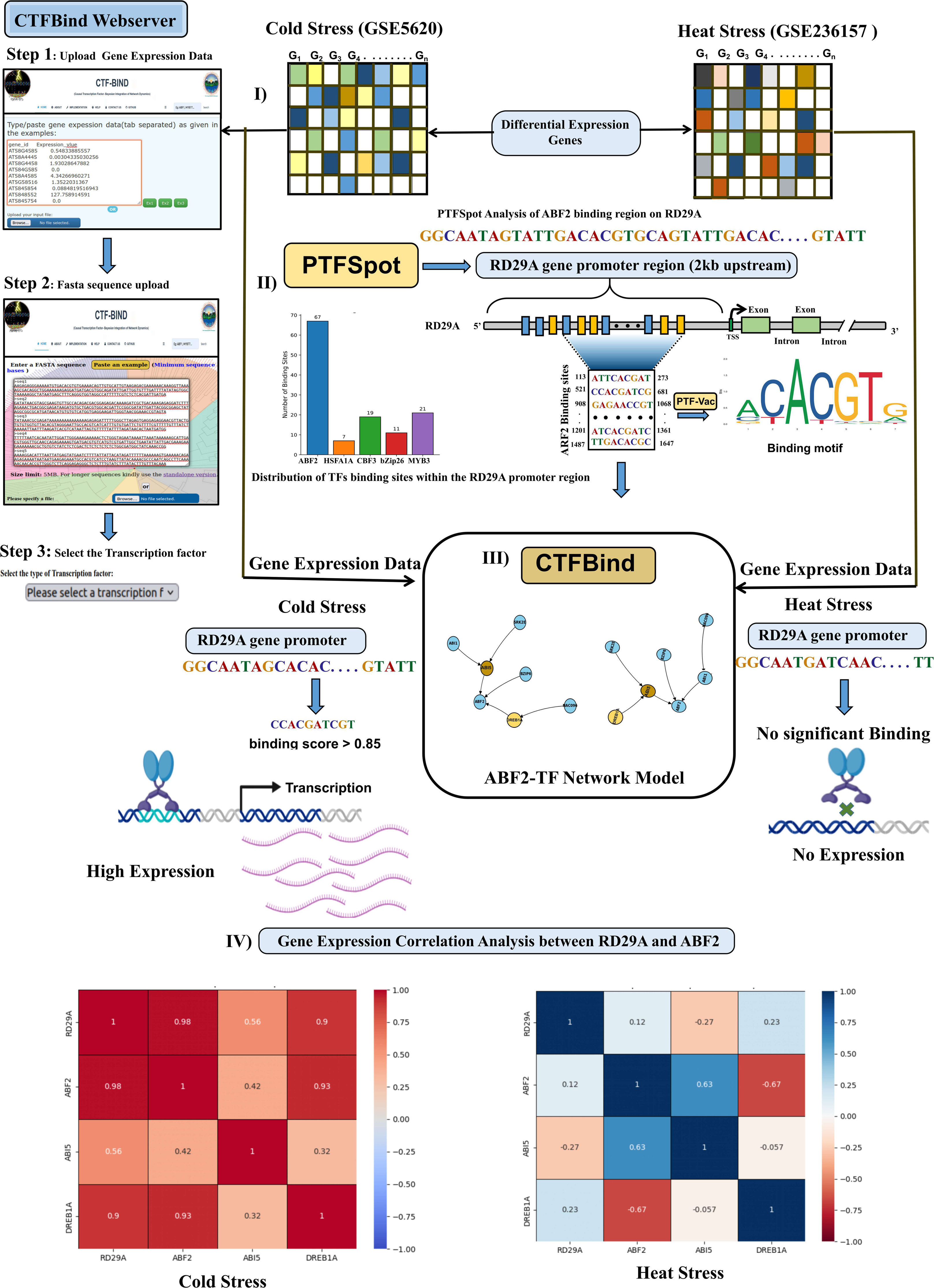
Application of CTF-BIND on RD29A gene as a case study with implementation of web-server.

### CTF-BIND: A modular platform for exploring Transcription Factor networks

CTF-BIND provides an intuitive interface for analyzing condition-specific TF networks, integrating six interconnected modules to explore regulatory dynamics under abiotic stresses. The “Quick Search” feature (top-right homepage) enables rapid access via four dropdowns: abiotic stress type, tissue, TF, and time-point. Users retrieve TF network metadata, including protein-protein interactions (PPIs), gene expression profiles, functional enrichment, pathway mappings, and TFBS visualizations.

#### Visualization and core modules

Networks are rendered using D3.js, where nodes (genes/TFs) are color-coded (active: colored; inactive: white) and edges denote regulatory relationships, with widths proportional to interaction weights. The platform’s six modules synergize as follows:

**1. Network Module:** Leverages NetworkX to compute topological metrics (degree distribution, centrality) and detect communities via greedy modularity optimization. Communities are visualized as force-directed graphs (D3.js), with shared colors indicating co-module membership. Network-wide properties (e.g., clustering coefficients) are tabulated.
**2. Interaction Module:** Distinguishes targeting (regulatory directionality) and binding (physical interactions) relationships. TFs (yellow squares) and target genes (purple circles) form interactive networks, elucidating transcriptional regulation hierarchies.
**3. Expression Module:** Displays RNA-seq-derived expression of TF-regulated genes as interactive bubble charts (Plotly). Bubble sizes reflect normalized expression levels, with axes representing genes (x) and expression values (y).
**4. Ontology Module:** Integrates TAIR-curated Gene Ontology (GO) terms, displaying biological processes, molecular functions, and cellular components upon node selection. Functional enrichment analysis via g:Profiler highlights top 20 GO terms, while pathway mappings (KEGG API) illustrate gene roles in cellular processes.
**5. Motif Visualization Module:** Renders TFBS sequences and locations using Feature Viewer (promoters: blue; exons: gray; binding sites: red). Sequence logos, generated via Logomaker, depict nucleotide conservation at binding sites.
**6. Comparison Module:** Enables side-by-side analysis of networks across conditions. Users select two stress contexts to view condition-specific D3.js networks, topological properties, and shared modules (greedy modularity), with edge widths reflecting interaction confidence.

A Flask backend bridges MongoDB (node properties, GO terms) and PyMongo, while Biopython interfaces with KEGG for pathway data. Interactive elements (Ajax, D3.js) ensure real-time updates, maintaining user engagement. CTF-BIND’s modular design balances analytical depth with usability, empowering researchers to dissect transcriptional regulation under dynamic stress conditions.

This case study demonstrates the utility of CTF-BIND-DB in dissecting the dynamic transcriptional regulatory network of HSFA1 (Heat Shock Transcription Factor A1) under heat stress in *Arabidopsis thaliana.* Heat shock-responsive genes, encoding proteins like heat shock proteins (HSPs), are crucial for maintaining cellular homeostasis under heat stress by preventing protein denaturation and aggregation (Liu et al., 2011; Ohama et al., 2017). Using CTF-BIND-DB, we investigated how HSFA1’s interactions shift dynamically across different heat stress time points (**Figure 11)**.

**Figure 11.**
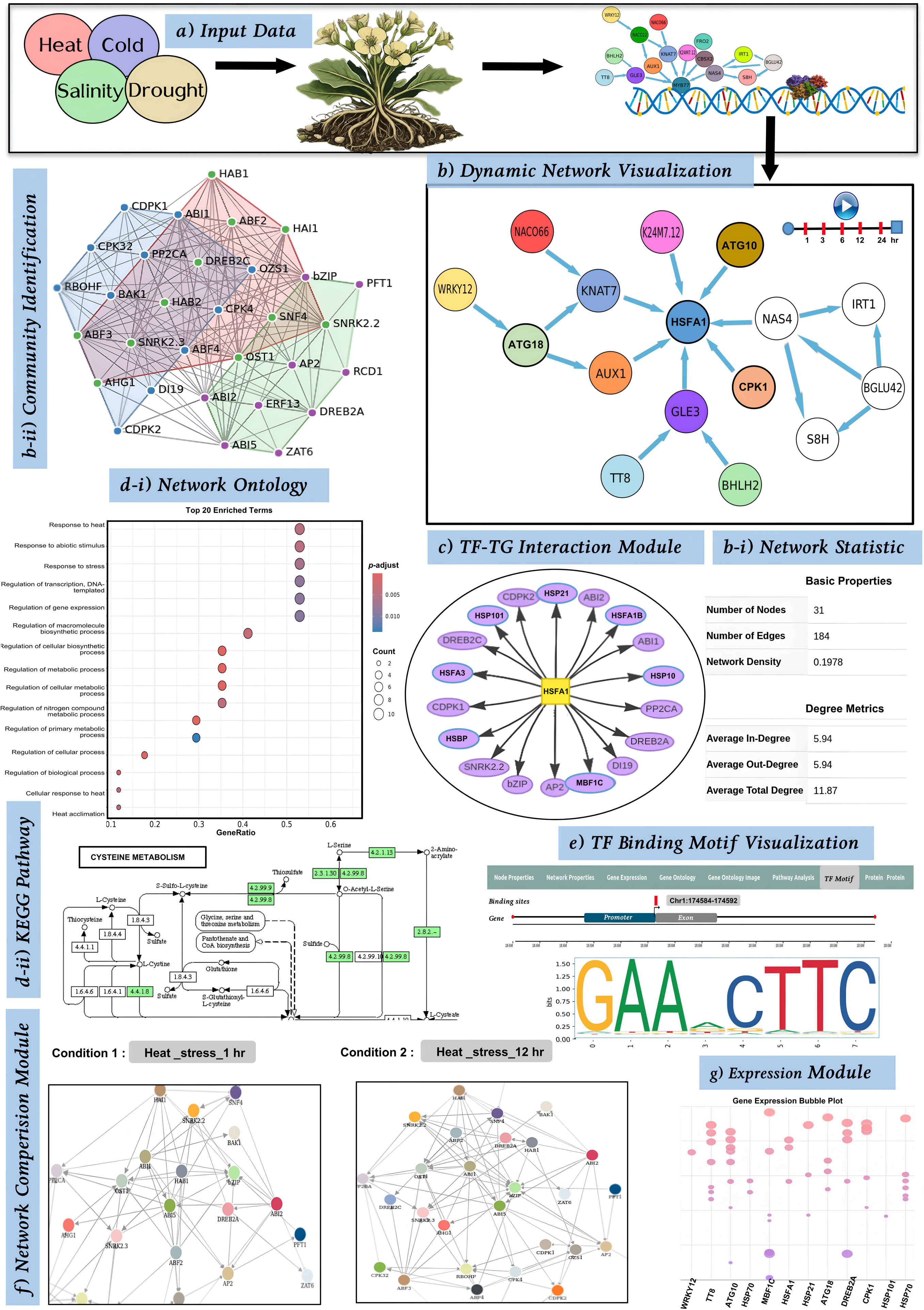
CTF-BIND-DB Implementation and Application. This figure demonstrates the application of CTF-BIND-DB to analyze the heat stress response of HSFA1 in *Arabidopsis thaliana*. **(a)** The Home Page provides a user interface for data input and stress selection. **(b)** Network Module visualizes the HSFA1 regulatory network, including network statistics (b-i) and community structures (b-ii). **(c)** Interaction Module shows specific HSFA1-target gene interactions. **(d)** Network Ontology Module reveals enriched GO terms (d-i) and KEGG pathways (d-ii) associated with HSFA1’s targets. **(e)** Motif Visualization Module highlights HSE motifs in target gene promoters. **(f)** Network Comparison Module compares the HSFA1 network under different heat stress time points (1h vs. 12h). **(g)** Gene Expression Module displays the expression dynamics of HSFA1 and its target genes under heat stress.

#### Network Analysis

HSFA1 regulates 47 heat shock-responsive genes (HSP70, HSP90, HSP101, sHSPs), exhibiting high degree (0.85), closeness (0.76), and betweenness centrality (0.71).

#### Interaction Module

HSFA1 binds directly to promoters of target genes, with strongest interaction seen for HSP70 (edge weight 0.92).

#### Expression Module

HSFA1 and its targets show early-stage expression peaks (0–3h), declining after 6h, aligning with transient stress responses.

#### Ontology Module

GO analysis reveals significant enrichment in “Response to heat” (GO:0009408, FDR = 0.0237) and KEGG pathways related to protein processing.

#### Motif Visualization

Identifies heat shock elements (HSEs) in target gene promoters, confirming HSFA1’s direct regulatory role.

#### Comparison Module

HSFA1 targets 47 genes under heat stress, but only 12 under cold stress, emphasizing stress-specific regulatory shifts.

This case study exemplifies the power of CTF-BIND-DB in dissecting complex regulatory networks under abiotic stress. By integrating diverse data types and providing user-friendly modules, CTF-BIND-DB facilitates a deeper understanding of TF function and stress response mechanisms. These findings provide valuable insights into plant stress biology and pave the way for developing strategies to enhance crop resilience to environmental challenges.

## Conclusion

In this study, causal networks for transcription factor (TF) binding were identified using diverse high-throughput data sources, experimental conditions, and cell lines. These networks captured a broader set of functional TF-target gene (TG) interactions, validated against existing experimental data. The approach not only re-validated the effectiveness of the networks but also demonstrated their ability to accurately predict TF binding profiles when used to train a deep learning system. The causal networks served as training modules, enabling predictions based solely on RNA-seq or transcriptome profiling data mapped to these networks. By exploring various causal paths, the system determines the TF binding profiles for specific experimental conditions.

The resulting software, CTF-BIND, revolutionizes TF binding profiling by eliminating the need for separate ChIP-seq experiments. It allows researchers to directly and accurately predict TF binding profiles from transcriptome data, making this process more accessible and scalable. Complementing this, CTF-BIND-DB (https://hichicob.ihbt.res.in/ctfbind/) was developed as a comprehensive database that captures dynamic shifts in regulatory pathways under diverse conditions. CTF-BIND-DB not only provides binding motifs and TF-TG interaction profiles but also includes detailed network ontology and functional information for downstream analysis. It facilitates the exploration of spatio-temporal dynamics of TF binding and their regulatory networks, offering researchers a centralized platform for integrating multi-omics data, performing functional annotations, and analyzing stress-responsive pathways.

Future studies should emphasize the relationship between TFs and TGs, especially under different conditions. The availability of additional high-throughput data, such as ChIP-seq and RNA-seq, will help expand the models to include more TFs and extend their applicability to other species. Incorporating DNA modifications, epitranscriptomics data, and protein-DNA interaction maps into CTF-BIND and CTF-BIND-DB could further enhance their utility, bringing us closer to comprehensive, system-level maps of TF biogenesis. These advancements will significantly contribute to understanding TF regulation, improving crop resilience, and driving innovations in plant stress biology.

## Acknowledgments

The work was carried out under the aegis of The Himalayan Centre for High-throughput Computational Biology (HiCHiCoB), a BIC supported by DBT, Govt. of India. UB is thankful to DBT, India for financial support as DBT-JRF. SG, AK, and AS are thankful to DBT, India for financial support as project associateship. UB and SG are also thankful to Academy of Scientific and Innovative Research (AcSIR) for their Ph.D. enrollment. All authors are thankful to the Director, CSIR-IHBT, for his kind support for this study. This MS has CSIR-IHBT MSID ##.

## Author’s contributions

UB carried out the major parts of this study. AS and SG developed the web-server of CTF-BIND. SG and AK assisted in the deep learning implementation. UK assisted in the Bayesian Network construction. RS conceptualized, designed, analyzed, and supervised the entire study. UB and RS wrote the MS.

## Software and Data availability

All the secondary data used in the present study were publicly available and their due references and sources have been provided in **Supplementary Table S1-2**. The software has also been made available as a webserver at https://hichicob.ihbt.res.in/ctfbind/.

