## Supplementary figures and images for "Decoding stress specific transcriptional regulation by causality aware Graph-Transformer deep learning"

### Supplementary Figure S1

#### Transcription Factors

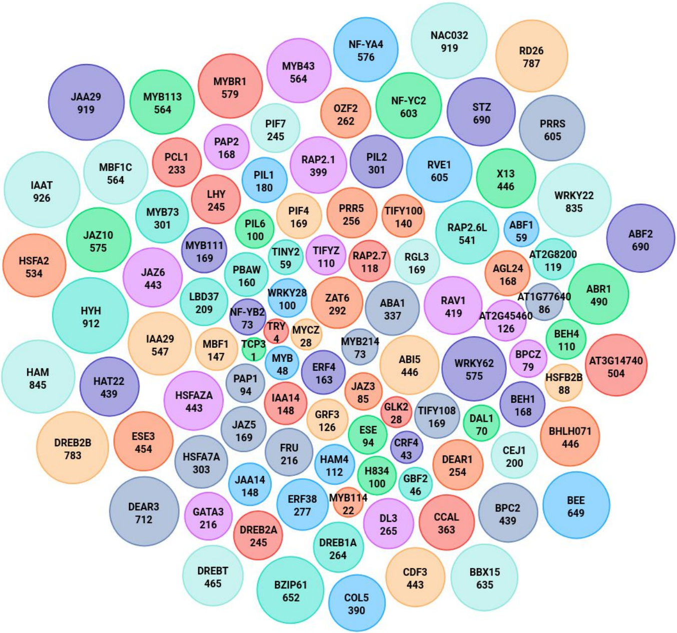

### Supplementary Figure S2

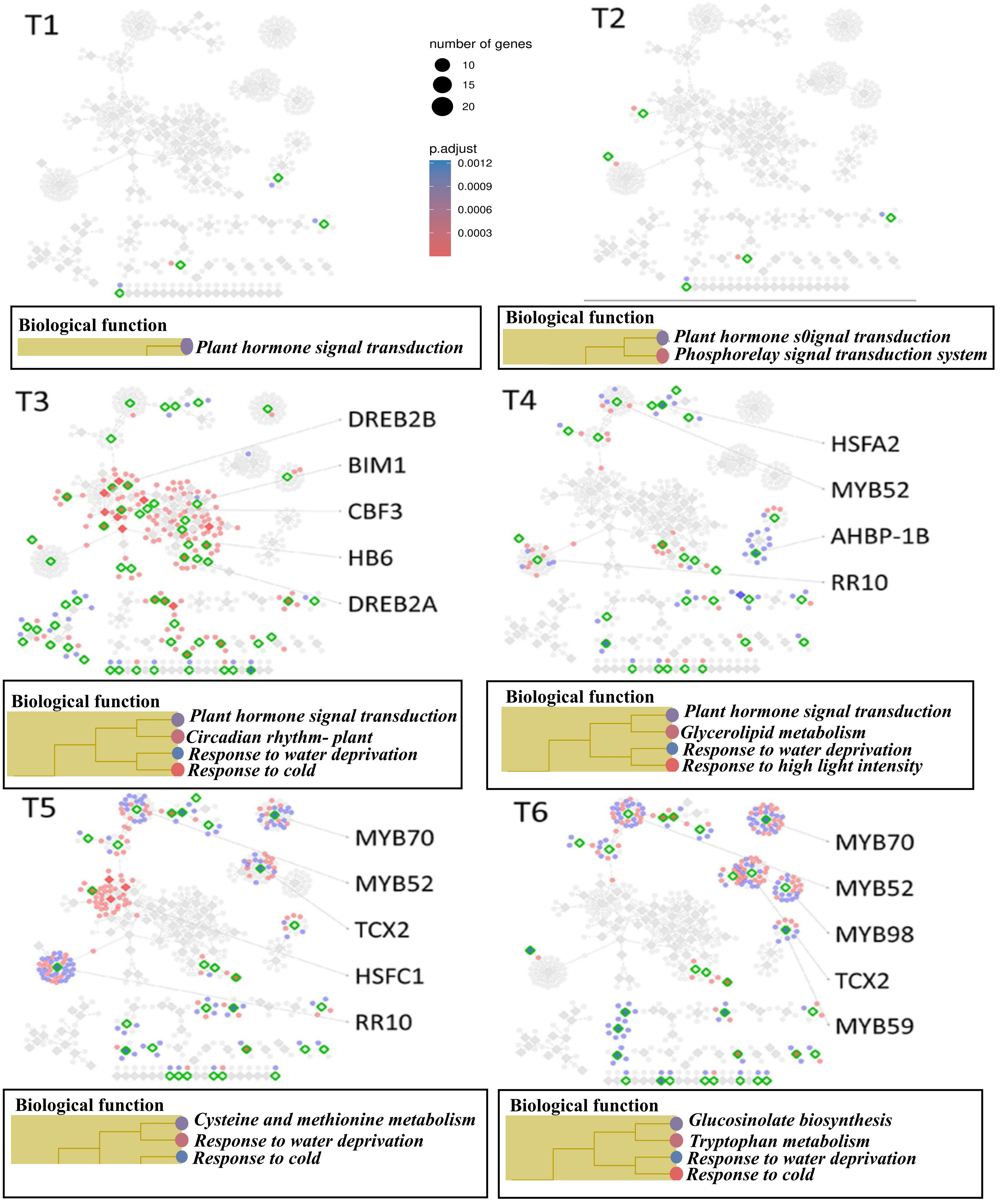

### Supplementary Figure S3

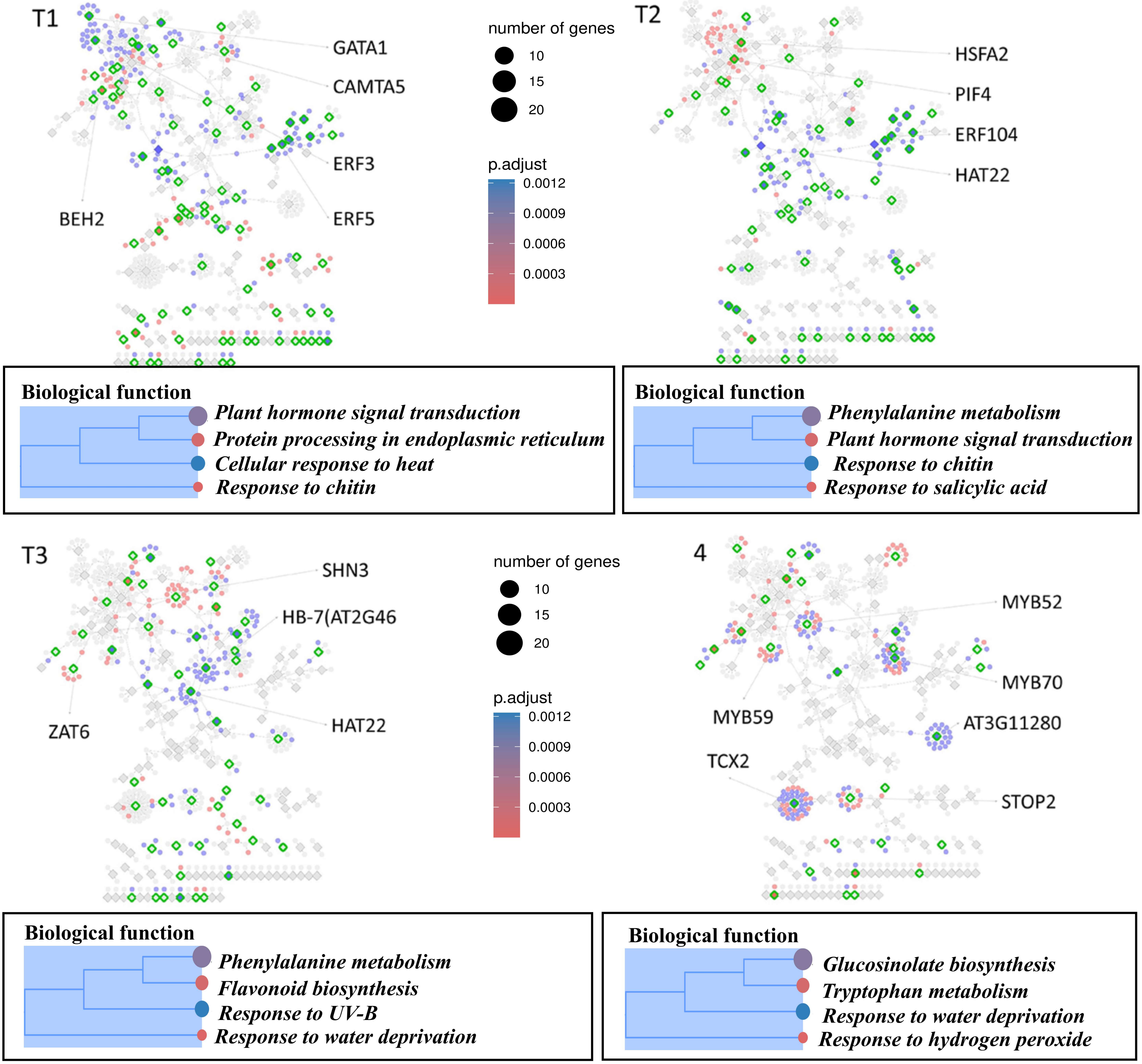

### Supplementary Figure S4

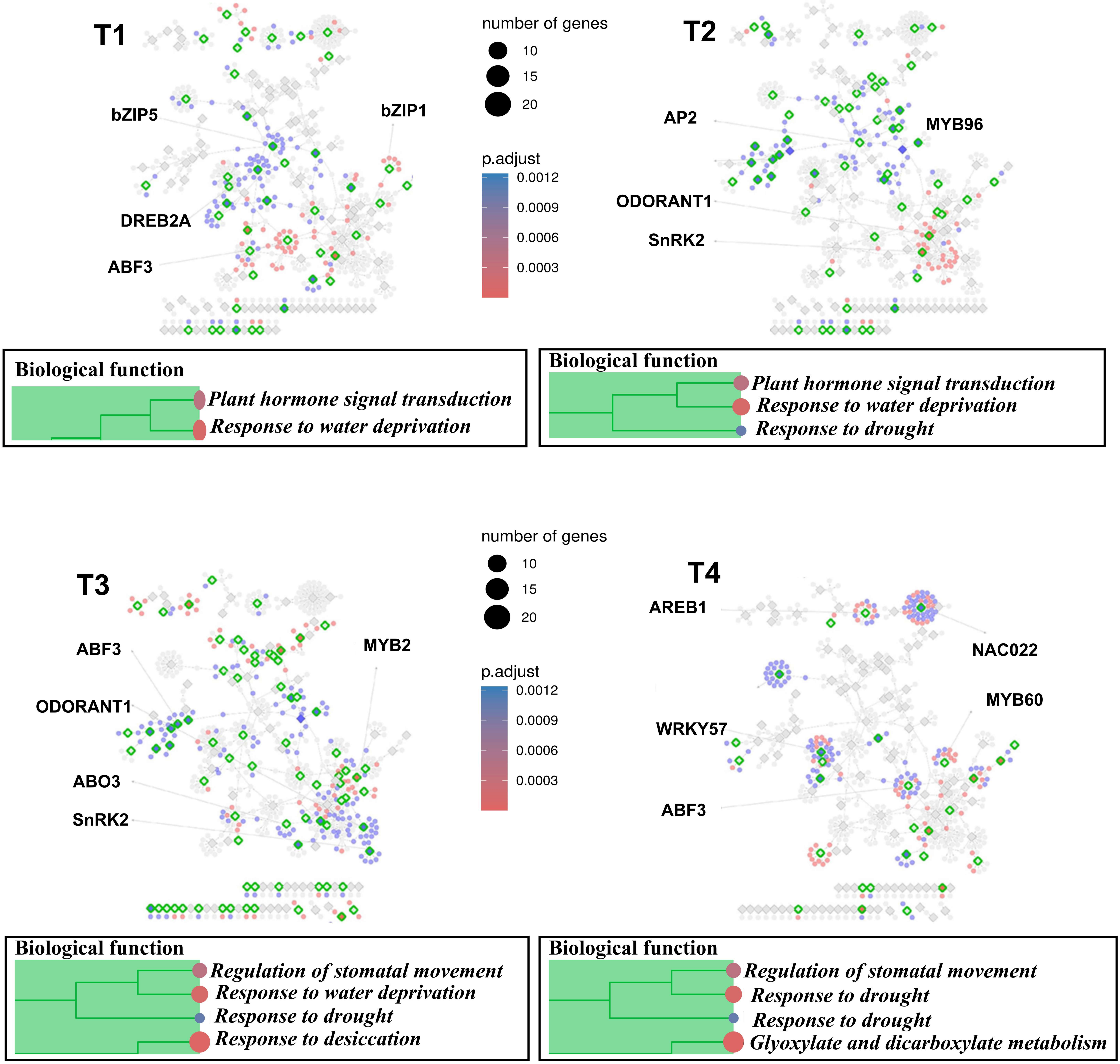

### Supplementary Figure S5

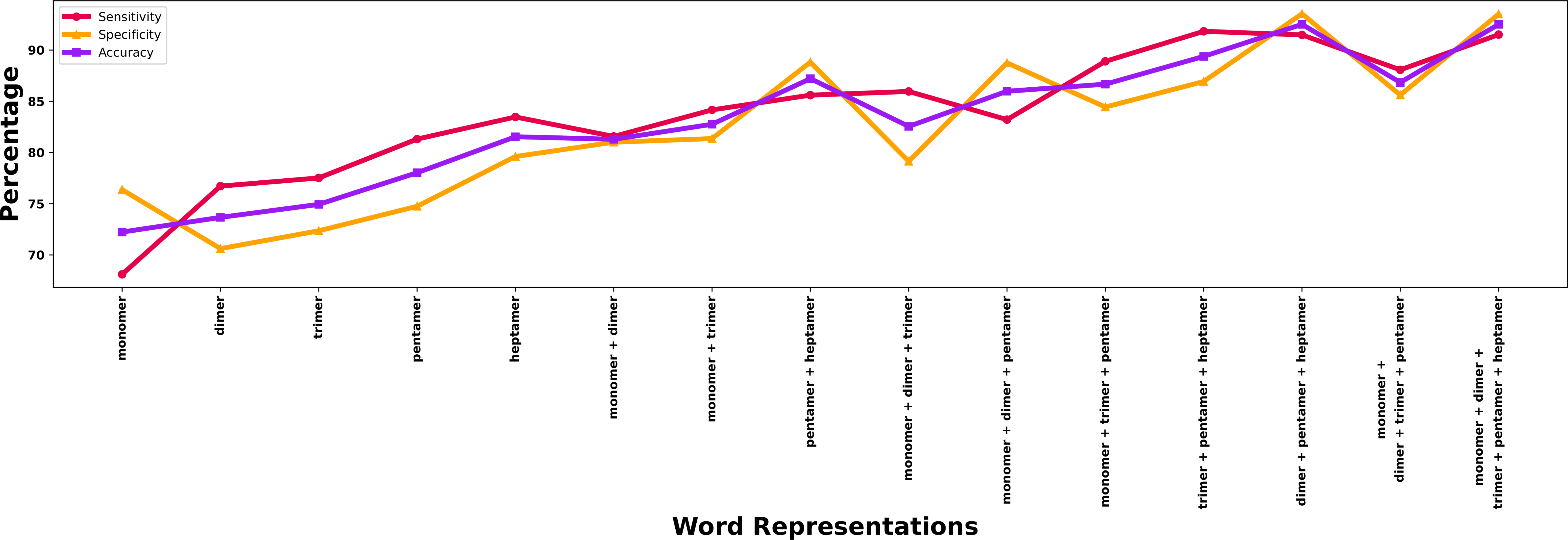

### Supplementary Figure S6

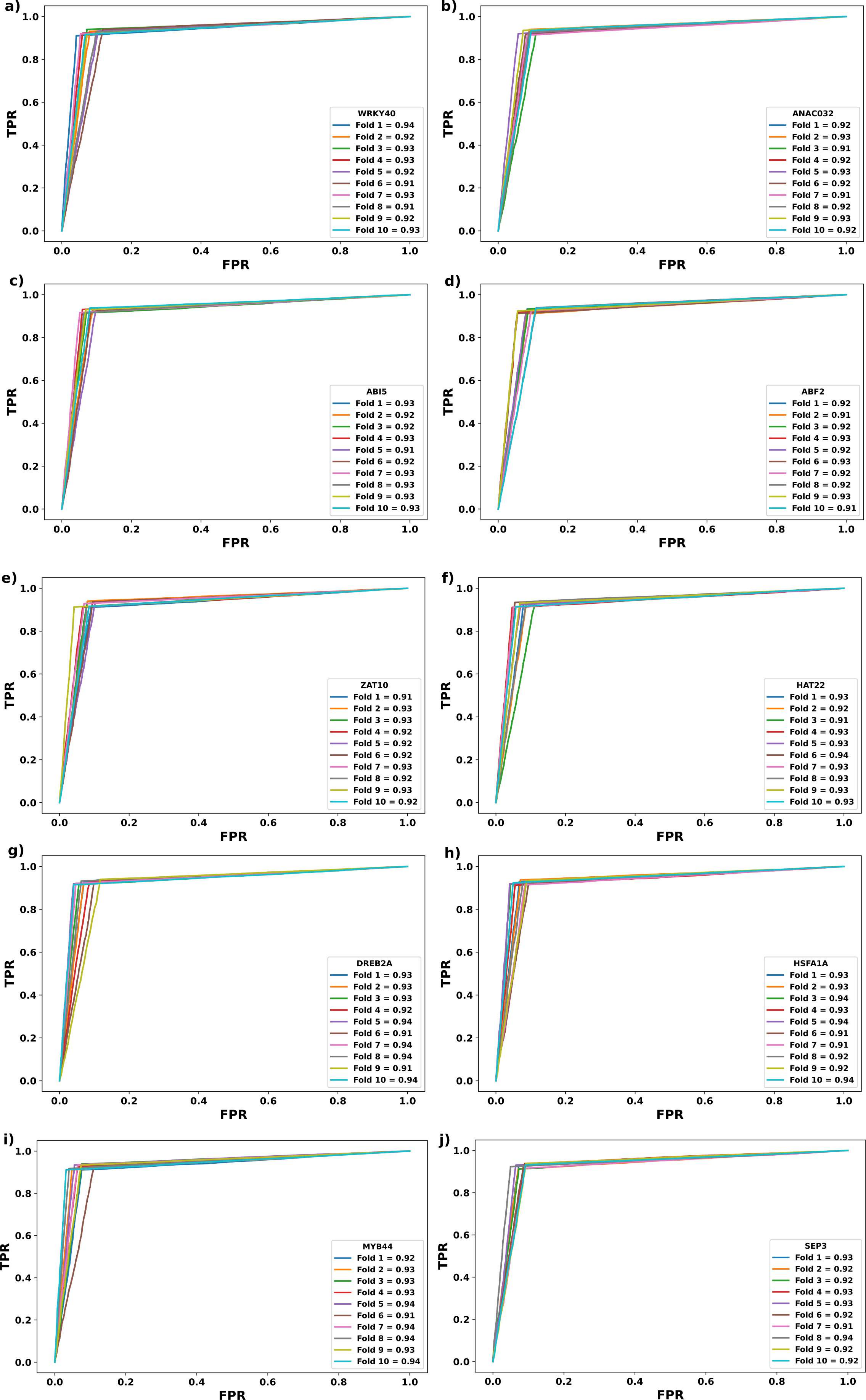

### Supplementary Figure S7

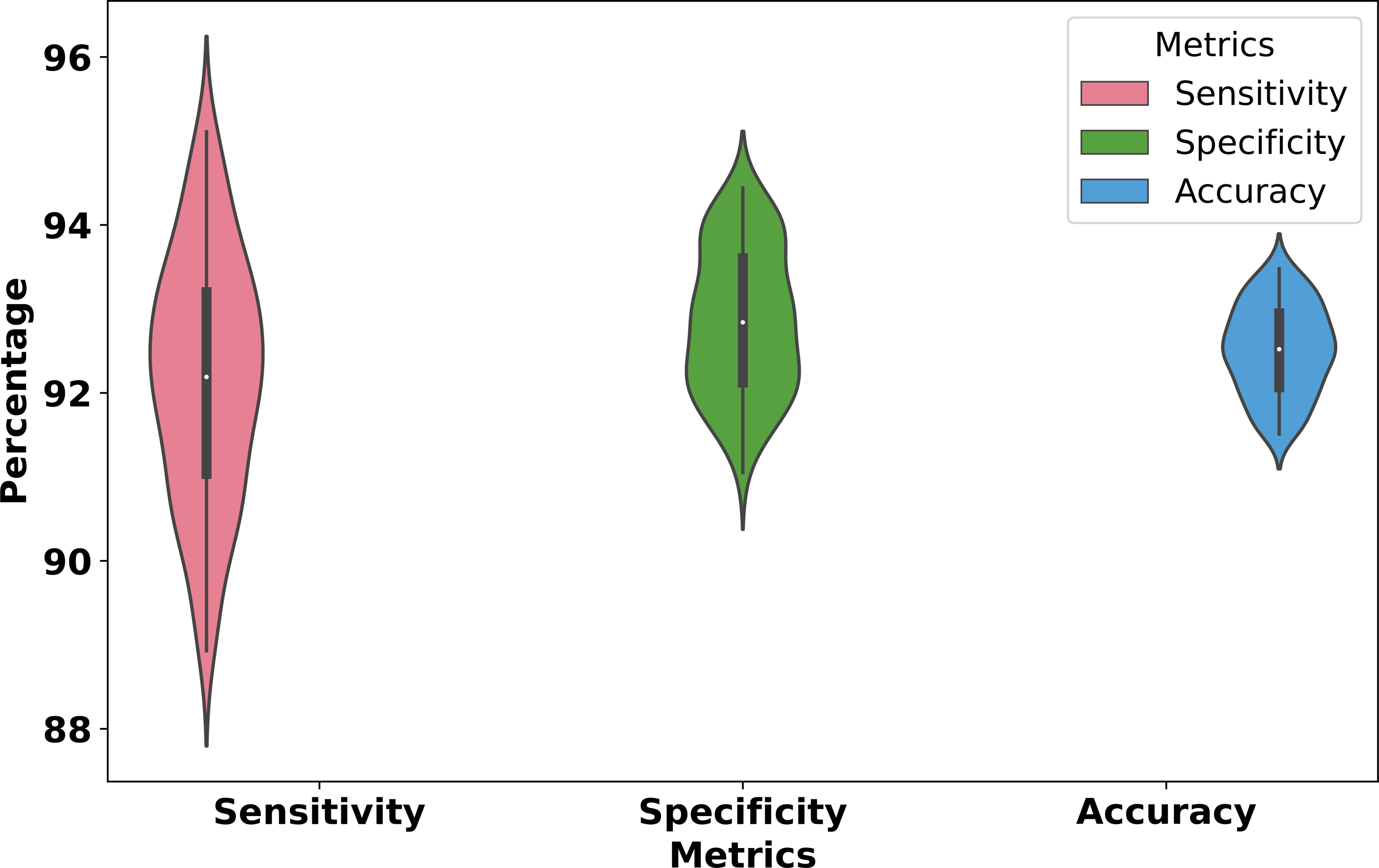

### Supplementary Figure S8

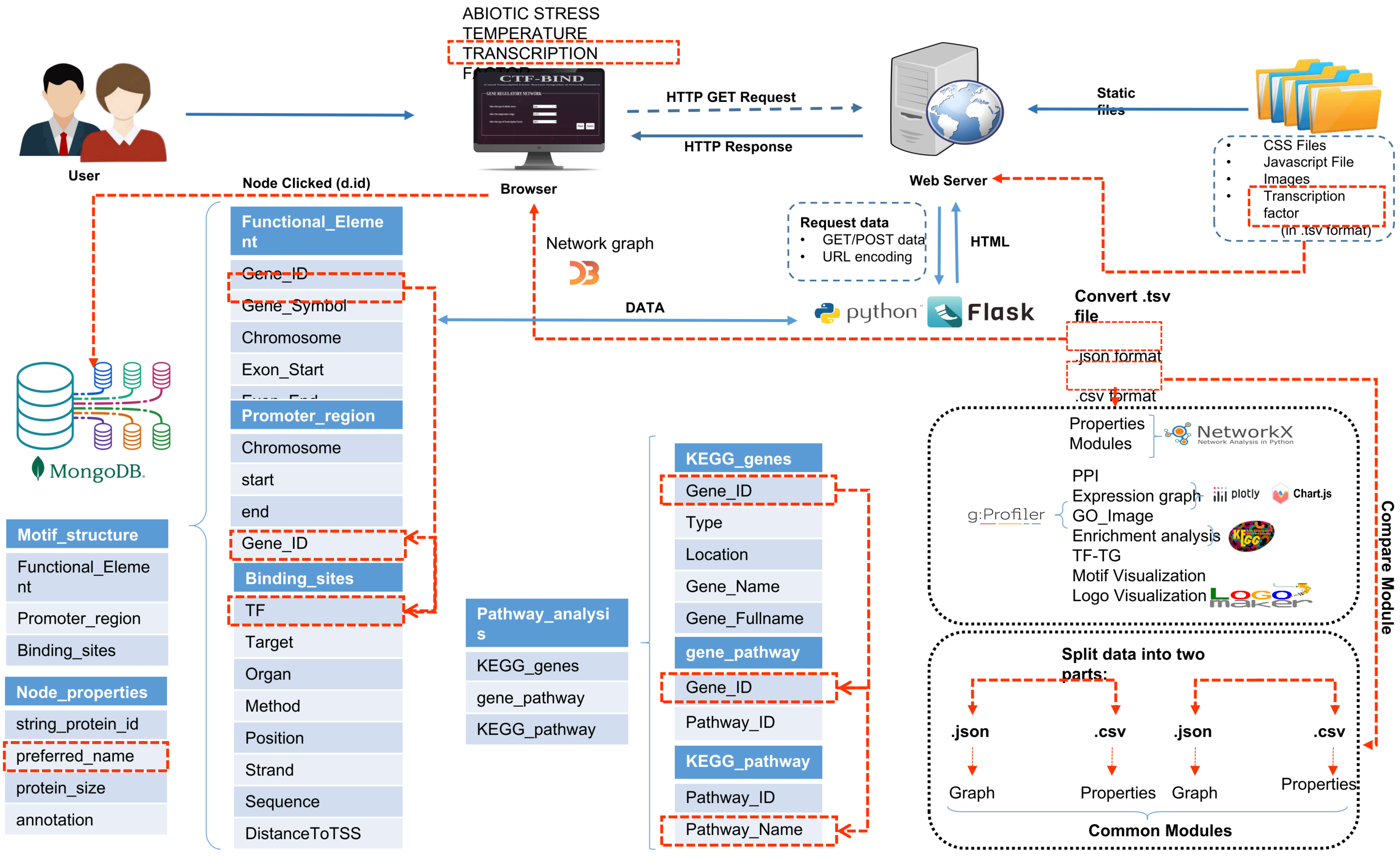
