## Supplementary material for "Decoding stress specific transcriptional regulation by causality aware Graph-Transformer deep learning": Supplimentary File 1

[1] Studio of Computational Biology & Bioinformatics,
The Himalayan Centre for High-throughput Computational Biology,
(HiCHiCoB, A BIC supported by DBT, India), Biotechnology Division,
CSIR-Institute of Himalayan Bioresource Technology (CSIR-IHBT),
Palampur (HP), 176061, India.

[2]Academy of Scientific and Innovative Research (AcSIR),
Ghaziabad, Uttar Pradesh- 201002

[3]ICAR-Indian Agricultural Statistics Research Institute, Library Avenue, Pusa, New Delhi, Delhi, India

Authors' email addresses:
UB:
AS:
AK:
SG:
UKP:
RS:

### Supplementary File 1:

### Materials and methods

#### Reconstruction of TF:TG associations using Bayesian network analysis (BNA)

ChIP-seq and RNA-seq data having same experimental conditions were collected from ChIP-Hub, GEO and ENA databases. Total 22 experimental conditions were common among all the two types of high-throughput data. Possible interacting partners were collected for each TF from various databse like STRING, AGRIS and stress knowledge map database (SKM) (Szklarczyk et al. 2019; Bleker et al. 2024). Maximum up to three steps of interactions were considered for the primary network construction. BNA was conducted separately for each experimental condition. The input data included gene expression levels, transcription factors (TFs), and associated genes derived from protein-protein interaction (PPI) data. TFs that had binding sites within the expressed genes under the specific experimental conditions were incorporated into the BNA. For any TF-TG interaction model, a TF was included if it had binding sites within the gene promoter regions. This analysis was repeated for each experimental condition, resulting in TF:TG associations for the expressed genes during the major steps of the process.

Given that TFs and their associated PPI components are key factors in TF actively binding, a more comprehensive approach was taken using structural equation modeling (SEM). This approach, previously applied successfully in our study investigating miRNA-RBP interactions (Pradhan et al., 2021), allows for the analysis of complex relationships within the regulatory network. SEM consists of two main components: the response (dependent variable) and the independent variables. The primary goal of SEM is to estimate the potential TFs involved at each step. As the data (expression levels of TFs and associated genes from PPI) were continuous, a multivariate gaussian distribution was assumed for all nodes throughout the study. The basic model used was a p-dimensional random vector ($X = (X_1, X_2, \ldots, X_p)$) with a joint distribution ($P = (X_1, X_2, \ldots, X_p)$). Here, ($X_1, X_2, \ldots, X_p$) represent the nodes in the network corresponding to TFs and associated genes from PPI.

Bayesian networks (BNs) are directed graphical models where the edges represent conditional independence constraints implied by the joint distribution of X:

$$P(x_1, \cdots, x_n) = \prod_{i=1}^{n} \theta(x_i | parents(X_i)) \quad (14.1)$$

$$P(a|b) = \frac{P(a \wedge b)}{P(b)} \quad (13.3)$$

$$\mathbf{P}(Y) = \sum_{z \in Z} P(Y, z) \quad (13.6)$$

**(1)**

where ($x_i$) denotes the parent set of ($X_i$) and theta represents the parameters defining the conditional probability distribution (CPD) for ($X_i$). The steps for constructing the TF model are as follows with their mathematical implementation:

1 **Estimation of Directed Acyclic Graphs (DAGs):** TFs with their respective PPI partners were modeled based on gene expression data and their associated proteins. The binding sites of TFs on genes were used as prior information. We used scored based learning such as Hill Climbing (HC) (Selman et al. 2006). it is a local search algorithm used to optimize a scoring function over the space of possible DAGs. In BNA structure learning, the HC algorithm iteratively improves a DAG structure by making local changes (such as adding, deleting, or reversing an edge) to maximize a scoring function, typically the Bayesian Information Criterion (BIC) score.

2 Algorithm 1: Hill Climbing for DAG Estimation

Input:

$D$ : Gene expression data.

$PPI$ : Protein-Protein Interaction data as prior knowledge.

$\lambda$ : Regularization parameter for incorporating PPI influence.

$G^{(0)}$ : Initial Directed Acyclic Graph (DAG), which could be based on prior knowledge. $MaxIter$ : Maximum number of iterations to perform.

2.1 Output:

2.2 $G^{(T)}$ : Estimated DAG that represents the gene regulatory network.

2.3 **Initialize:**

2.4 Set $t=0$

2.5 Initialize the current graph $G^{(t)}=G^{(0)}$

2.6 Compute the initial score $S_{PPI}\left(G^{(t)}|D\right)$

2.7 **Repeat until convergence or until the maximum number of iterations is reached:**

2.8 Set $best_{score}=S_{PPI}\left(G^{(t)}|D\right)$

2.9 Set improved= *False*

2.10 **For each possible edge modification (add, delete, or reverse) that maintains the DAG property:**

2.11 Generate a neighboring graph $G^{'}$

2.12 Compute the score for this new graph:

2.13 $$S_{PPI}\left(G^{(t)}|D\right)=logP\left(D|G^{'}\right)-\frac{|G^{'}|}{2}logN+\lambda\sum_{(i,j\in E^{'})}PPI(i,j)$$

2.14 where:

2.15 $P\left(D|G^{'}\right)$ is the likelihood of the data given the graph $G^{'}$

2.16 $|G^{'}|$ is the number of edges in the graph $G^{'}$

2.17 $N$ is the number of samples in the gene expression data.

2.18 $\sum_{(i,j\in E^{'})}PPI(i,j)$ is the sum of PPI scores for all edges in the graph $G^{'}$

2.19 **If** $S_{PPI}\left(G^{(t)}|D\right)>best_{score}$ :

2.20 Update the graph: $G^{(t+1)}=G^{'}$

2.21 Update the best score: $best_{score}=S_{PPI}\left(G^{(t)}|D\right)$

2.22 Set improved = *True*

2.23 **If** improved = *True* :

2.24 Update the current graph: $G^{t}=G^{(t+1)}$

2.25 Increment the iteration count: $t = t + 1$

2.26 **Else** (if no improvement was found):

2.27 Stop, convergence has been reached

2.28 **Return** the estimated DAG: $G^{(T)} = G^{(t)}$

**2.** Identification of significant DAGs between TFs and TGs, TFs with their respective PPI partners were taken out from the total DAGs considering a suitable a statistical test (e.g., chi-square test) was used.

**3**. It was assumed throughout the study that the data were generated from a multivariate gaussian distribution, where error covariance matrix is a positive definite. Thus, the significant DAGs obtained in the previous step were directly modeled through a generalized linear model.

$$\mathrm{X} = \mathbf{B}^{\mathrm{T}}\mathrm{X} + \mathrm{E}$$

This is called a SEM for total observations **X**. **B** is the weighted adjacency matrix of a directed graph and E∼ N (0, $w_j^2$).

**4.** The significant DAGs were modeled using SEM. To avoid the high-dimensional data structure (i.e. n<<p), sparse regularized penalty was used. Following LASSO model was considered for the data:

$$\min_{\beta}\left(\sum_{i=1}^{N}\left(Y_i - \beta_{0i} - \sum_{j\in \mathrm{pa}(i)} \beta_{ij} Y_j\right)^2 + \lambda \sum_{i,j} |\beta_{ij}|\right) \quad \textbf{(2)}$$

where: $\lambda$ is a regularization parameter controlling the sparsity of the network.

5. Once the network structure is learned, the next step is to estimate the parameters (e.g., edge weights, regression coefficients) Maximum Likelihood Estimation (MLE): Used to estimate parameters that maximize the likelihood of observing the given data under the assumed model.

$$(D|\mathrm{G},\theta)$$

where $\theta$ represents the parameters of the model (e.g., regression coefficients $\beta_{ij}$ for SEM).

**6.** Further, to improve the estimation of parameter, weighted adjacent matrix was represented by B as B = est($\beta$) which was used to estimate the conditional variance by the given formula:

$$\mathrm{Est}(W_j)^2 = \mathrm{var}(x_j - X(\mathrm{est}(\beta_j))) \quad \textbf{(3)}$$

$\Omega = \mathrm{diag}(\mathrm{est}(W_1)^2,\ldots,\mathrm{est}(W_p)^2)$ was applied as a variance matrix and combining [est($B$),est($\Omega$)] to calculate the variance covariance matrix Σ and accuracy for each parameter. A suitable convergence criteria (Error tolerance < $10^{-4}$), precision value > = 85% and an alpha threshold of 0.05 were considered for the selection of the parameters. These parameters decided how larger the effect size (positive/negative) was between TGs and TFs.

The algorithm followed in BNA was performed using "bnlearn", and "ccdr Algorithm", both are modules of R (Scutari et al. 2009). The associated basic steps followed in the current approach are described in **Figure 2**.

#### Optimization of the sequence transformer system

The sequence transformer system was optimized using Bayesian optimization. This transformer encoders had a multi-head attention layer, where 14 self-attention heads were found to work best. The input sequence was padded if the length was shorter or longer than 160 bases to ensure a constant size of the input matrix. The output from the multi-head attention layer was passed into the dropout layer with a dropout fraction of 0.1. Later, this result was normalized by another layer called the normalization layer. This layer was followed by a third layer called feed-forward layer with 38 nodes, followed by another dropout layer with a dropout fraction of 0.2. A feed-forward layer with 12 nodes, followed by another normalization layer, followed the above one. Different activation functions for the feed forward layer layers were explored among the available activation functions. Finally, two dense layers were configured, with SELU and RELU activation functions, respectively. Afterwards, the result of the normalization layer was passed to the GlobalAveragePooling1D layer. A binary cross-entropy loss function was used to calculate the loss and "Adam" optimizer was used to adjust the weights and learning rates. The learning rate of the optimizer was set to 0.015, and the model was trained using 45 epochs and batch size of 32. Since the problem in this study was not translation but classification, decoders were not needed. The number of encoders and performance was investigated, and it was found that increasing the encoder layers did show a significantly change in efficiency and only decreased after the ninth encoder layer. Thus, we went with eight encoder layer. This sequence transformer part output is flattened and concatenated with the Graph transformer output, which was then passed into the second transformer encoder block which derived the hidden features and their relationships which got structured, on which classification could be done in much superior manner. The related information about optimization towards the final model is listed in Supplementary Table 3 Sheet 1.

#### Graph representation for the graph transformer

The Graph Transformer architecture integrates three primary data components: graph structure, node features, and node-level metrics derived from TF-specific causal networks under 46 distinct abiotic stress conditions (drought, cold, heat, and salinity). These networks are conceptualized as Protein-Protein Interaction (PPI) graphs, with each graph corresponding to a specific TF under a given condition. The mathematical formulation for each component is detailed below.

**Graph structure**

The TF-specific causal network for condition **"c"** and transcription factor $TF_i$ is represented as a directed graph $G_{c,TF_i} = (V_{c,TF_i}, E_{c,TF_i})$, where: $V_{c,TF_i}$represents the node set (proteins, including TF and target genes),$E_{c,TF_i}$denotes the directed edge set, where (u,v) $\in$ $E_{c,TF_i}$ indicates a regulatory interaction from protein **"u"** to **"v"**. The graph structure is encoded in an adjacency matrix $A_{c,TF_i} \in \mathbb{R}^{N_{c,TF_i} \times N_{c,TF_i}}$, where $N_{c,TF_i} = |V_{c,TF_i}|$ is the number of nodes in the graph. Matrix entries are defined as:

$$A_{c,TF_i}[u,v] = \begin{cases} 1, if (u \to v) \in E_{c,TF_i} \\ 0, otherwise \end{cases} \quad \textbf{(4)}$$

The directed nature of these graphs ensures proper representation of regulatory relationships while maintaining acyclicity to conform with causal assumptions.

**Node features**

Each node $v \in V_{c,TF_i}$ is characterized by a feature vector $x_v$ derived from condition-specific gene expression data. The collective node features form a feature matrix $X_{c,TF_i} \in \mathbb{R}^{N_{c,TF_i} \times F}$, where F represents the z-score normalized gene expression value.

**Node-Level metrics**

Additionally, we calculated five other metrics $M_v$ using the NetworkX Python package (Hagberg et al. 2008) to quantify the centrality or importance of nodes within the network to capture the structural properties of each network. These consist of degree, betweenness, eigenvector, and closeness centrality, and the PageRank score. This comprehensive representation ($A_{c,TF_i}$, $X_{c,TF_i}$, $M_v$) captures both structural and functional characteristics of the TF-specific causal networks, enabling effective learning of local and global regulatory patterns by the Graph Transformer.

#### Graph Transformer architecture for condition-specific TFBS identification

The Graph Transformer learns features from the TF causal network, which represents the interactions between transcription factors (TFs) and their target genes. The nodes represent TFs or target genes, and edges represent regulatory relationships.

**Node Embedding Initialization**

Each node in the TF causal network is initialized with a feature vector $h_v^0$, which combines:

Each node $v$ in the TF causal network is initialized with a feature vector $h_v^0$, which integrates three critical components to represent the node's biological and topological properties. The Node Features

$x_v$ capture biological characteristics of the node, such as gene expression levels, providing context about the gene or transcription factor's functional state. The Centrality Encoding $M_v$ highlights the node's importance within the network, incorporating metrics like degree, betweenness, or closeness centrality to reflect its role in regulatory interactions. These components are concatenated to form the initial representation as $h_v^0 = concat(x_v, M_v)$, ensuring that each node's initialization encodes both biological relevance and network topology.

**Edge embedding initialization**

For each edge $(u, v)$in the TF causal network, an edge embedding $e_{uv}^{(0)}$is initialized to encode the type of regulatory interaction between the connected nodes. This is achieved using $e_{uv}^{(0)} = Embed(EdgeType(u, v))$, where $EdgeType(u, v)$ specifies the nature of the interaction, such as activation or repression. The learnable embedding function $Embed(.)$maps these interaction types into a continuous vector space, enabling the model to distinguish between different regulatory roles and incorporate this information during attention calculations.

**Attention mechanism**

The Graph Transformer employs a scaled dot-product attention mechanism to model the relationships between nodes and their neighbors in the TF causal network. The attention score between a pair of nodes **"u"** and **"v"** is computed as:

$$\alpha_{uv} = \frac{exp\left(LeakyReLU\left(a^T[W.h_u \vee W.h_v]\right) + \phi(u,v)\right)}{\sum_{k \in N(v)} exp} \quad \textbf{(5)}$$

where, W is a learnable weight matrix used to transform node embeddings, **a** is a learnable vector used for scoring attention, $[\cdot \vee \cdot]$ denotes concatenation of two vectors, $\phi(u, v)$ represents the spatial encoding (e.g., shortest path distance or a predefined measure of relatedness between nodes u and v) , $N(v)$ denotes the set of neighbors of node v.

**Node representation update**

The node embedding at layer $l+1$ is updated as:

$$h_v^{(l+1)} = \sigma\left(\sum_{u\in N(v)} \alpha_{uv}\cdot\left(w.\, h_v^{(l)} + e_{uv}^{(l)}\right)\right) \qquad \textbf{(6)}$$

where: $\sigma$ is a nonlinear activation function (e.g., ReLU), $\alpha_{uv}$ is the attention coefficient, $h_v^{(l)}$ is the node embedding of u at layer l, $e_{uv}^{(l)}$ is the edge embedding at layer l .

**Edge representation update**

The edge embedding $e_{uv}$ is updated at each layer to encode dynamic interactions as:

$$e_{uv}^{(l+1)} = W_e . e_{uv}^{(l)} + \sum_{p=1}^{P} W_p . Embed\left(Path_p|(u,v)\right) \qquad \textbf{(7)}$$

where: $W_e$ is a learnable transformation matrix for edge embeddings, $Path_p(u,v)$ encodes the *p-th* shortest path between nodes u and v, $Embed$(.) maps the path into a vector representation, $W_p$ is a learnable weight matrix for the *p-th* path.

This mechanism enables the Graph Transformer to dynamically refine both node and edge embeddings, capturing complex regulatory patterns and higher-order relationships in the TF causal network. It effectively models both local interactions and global dependencies, critical for analyzing transcription factor binding and regulatory influence. We implemented the graph-transformer with PyTorch geometric and adopted the “Adam” optimizer to optimize the parameters. Each node in the TF network was represented with a 160-dimensional learnable embedding vector, which was then fed into Graph Transformer layers that was 12 in number, to generate the network representations. Specifically, we utilized eight attention heads, each contributing to 20-dimensional latent representations for every Graph Transformer layer. The shortest path distance bound N was set to 3 for edge encoding between node pairs. This mechanism enabled the Graph Transformer to dynamically refine both node and edge embeddings, capturing complex regulatory patterns and higher-order relationships in the TF causal network. It effectively modeled both local interactions and global dependencies, which are critical for analyzing transcription factor binding and regulatory influence. To improve the stability and effectiveness of the learning process, skip connections and layer normalization were applied after each update to the node and edge representations. This approach helps address common issues in deep graph networks, such as over-smoothing and vanishing gradients. After down-sampling, the graph-transformer output is flattened and concatenated with the sequence transformer output for classification, and then passed into the second transformer encoder block. For multi-headed attention, the process is repeated based on the number of heads, and the individual attention score vectors are concatenated before being forwarded to the feed-forward network of the transformer encoder for further processing. The output from the multi-head attention layer is passed through a dropout layer to reduce over-fitting, followed by a layer of normalization. This output is then fed into a feed-forward layer, which sends its result to the next dropout layer in the stack. The feed-forward layer receives its input from the previous layer and is followed by a GlobalAveragePooling1D layer, which then passes its output to the third dropout layer. Various activation functions were tested for the layers using a pool of available functions. The output from the fourth dropout layer is passed through a LeakyReLU activation function, leading to a single-node classification layer. Binary cross-entropy is used as the loss function to calculate the error. The "Adam" optimizer adjusts the weights and learning rates, adapting learning rates based on a moving window of gradient updates, rather than accumulating all past gradients.

The transformer network extracts and structures hidden features and their relationships, making classification more effective. Since the present problem in this study was not translation but classification, decoders were not needed and instead the encoder output were taken as input for next step of extreme gradient boosting. For this purpose the output of the transformer was passed to the XGBoost classification part. XGBoost was the choice as it has consistently scored highest in

Kaggle benchmarking studies and performs exceptionally good on structured and curated features sets. **Figure 3** provides a snapshot of how this entire system is working.

#### Optimization of the GraphTransformer-Transformer-XGBoost system

Optimization of the hyper-parameters is an important step to derive the best possible model. To optimize the GraphTransformer architecture for condition specific transcription factor binding site (TFBS) identification, we employed a Bayesian optimization approach using Optuna. This allowed for efficient exploration of a defined hyperparameter search space to maximize model performance. The search space encompassed crucial architectural and training parameters, including the node embedding dimension (80-320), number of Graph Transformer layers (4-16), number of attention heads (4-14), shortest path distance bound (2-5), dropout rates (0.1-0.5), learning rate (1e-5 to 1e-3), batch size (16-128), and activation functions (ReLU, LeakyReLU, SELU, GELU). Optuna's Tree-structured Parzen Estimator (TPE) sampler guided the search, balancing exploration of new configurations with exploitation of promising regions, while a pruning mechanism terminated unpromising trials early to conserve computational resources. Optuna's TPE sampler efficiently navigated the hyperparameter space, dynamically adjusting the search based on observed performance. This approach allowed us to efficiently identify a high-performing configuration without exhaustively testing all possible combinations. The optimal configuration found included a node embedding dimension of 160, 12 Graph Transformer layers, 8 attention heads, a shortest path distance bound of 3, dropout of 0.15, a learning rate of 0.0005, a batch size of 32, and the LeakyReLU activation function. The optimized Graph Transformer was trained using the Adam optimizer with a learning rate scheduler, 50 epochs, and binary cross-entropy loss. To ensure the robustness of the results, 10-fold cross-validation was employed. This rigorous optimization strategy, combined with careful training procedures, resulted in a highly effective model for condition specific TFBS identification.

In the classification part of the hybrid Graph Transformer-Transformer-XGBoost, XGBoost takes input from the second fully connected layer of the second Transformers stack. Grid search was applied for hyperparameter optimization using scikit-learn function RandomizedSearchCV. Following hyperparameters were finalized after the grid search: params = {"eta/learning rate": 0.2, "max_depth": 6, "objective": "binary:logistic", "silent": 1, "base_score": np.mean(yt), "gamma": 7.81, "subsample": 0.85, "eta": 0.2, "colsample_bytree": 0.85, "n_estimators": 1400, "min_child_weight": 4.59, "eval_metric": "logloss", "reg_alpha": 147.2154, "reg_lambda": 0.0125, "tree_method": 'approx'}. Gradient boosted decision trees learn very quickly and may overfit. To overcome over-fitting shrinkage was used which slows down the learning rate of gradient boosting models. Size of the decision tree were run on different combinations of max-depth. Values changed until stability was gained as the logloss got stabilized and did not change thereafter. The final max_depth value was 6. The output from the XGBoost returned the probability score for each input sequence. The probability score indicated the confidence of each instance as TFBS or not. The final hyperparameters set for the output layer of the implemented model was: {“Activation function”: LeakyReLu, “Loss function”: binary crossentropy, “Optimizer”: Adam}. The related information about optimization towards the final model is listed in **Supplementary Table Sheet 1-3**.

The final model obtained was saved in “pt” format. Since the entire system is implemented here using Pytorch and scikit-learn, the “pt” format provided the model graph definition and weights to the pytorch structure while saving the model for classification purpose. Each and every hyper-parameter values involved to finalize this hybrid model were fixed using an in-house developed script which tested various combinations of values of the hyper-parameters to pick the best ones. This entire optimization process was done using two different approaches: Random search optimization and Bayesian optimizations. Figure 3 shows the detailed workflow of the implemented architecture.

#### Implementation of the webserver

##### Front-end Development

The implementation of CTF-BIND is divided into front-end and back-end. The front end of the platform was created using JavaScript (https://www.javascript.com/) and jQuery (https://jquery.com/), incorporating multiple visualization libraries: Plotly.js package (https://plotly.com/python/) and Chart.js (https://www.chartjs.org/) for data visualization, D3.js (https://d3js.org/) for network visualization, Logomaker Python package (https://logomaker.readthedocs.io/en/latest/) for motif sequence visualization, KEGG REST API (https://www.kegg.jp/kegg/rest/) for pathway analysis, and gprofiler2 REST API (https://biit.cs.ut.ee/gprofiler/page/r) for functional enrichment analysis (**Supplementary Figure S8**).

##### Back-end infrastructure

The server-side architecture utilized MongoDB (https://www.mongodb.com/) for data storage, Python (https://www.python.org/) for computational processing, and Flask (http://flask.pocoo.org/) for API development of CTF-BIND is being deployed on the Ubuntu Linux system (version 18.04.6 LTS) with the Apache HTTP Server for open and stable service. The database interactions were managed through PyMongo with asynchronous data retrieval via Ajax (**Supplementary Figure S8**).

##### Usage

The “Quick Search” feature on the top right section of the homepage provides an easy survey of the abiotic stress specific TF. The interface included four drop-downs for the user to select abiotic stress, tissue type, transcription factor, and time-point. Users can obtain basic information about the TF network, network modules, PPI, gene expression, GO, network functional enrichment analysis, pathway analysis, visualization of the TFBS and its sequence, TF-target gene network, and comparison of TF networks under various conditions.

CTF-BIND utilizes the D3.js library to visualize the gene regulatory network. In the network graph, each node represents a gene connected through an edge that defines the direction and nature of the relationship between the nodes. The width of an edge indicates the weight or value at which one node binds to another node. Each node is given a unique color to differentiate it from the others. The colored node reflects the active gene, as it is predicted to regulate the TF in the given condition. As the condition changes, the network graph also changes. Under a given condition, if a node loses its color (turns white), it reflects the non-functionality or inactivity of the gene in regulating the TF under that specific condition. Clicking on a node displays its properties and is stored in the MongoDB.

CTF-BIND offers the following six distinct modules for users to easily explore the TF network: network, interaction, expression, motif visualization, ontology, and comparison modules.

**Network module:** CTF-BIND studies the structure and dynamics of TF networks using the Networkx Python package (https://networkx.org/) (Hagberg et al., 2008). The properties of the overall network are displayed that consists of multiple attributes.

The study of topologies, such as degree distribution, clustering coefficient, and centrality measures, can provide insight into the organization, functions, biological processes, and important components of the network. Community detection within networks employs the greedy modularity optimization algorithm implemented in Networkx. The visually interactive representation of the modules is displayed below the table, which was built using D3js (force-directed graph). Nodes within the same module are represented by the same color.

**Interaction module:**

**Protein-Protein interaction (PPI):** The system analyzes two types of protein relationships.

1 Targeting relationships: Identifying nodes that target other network components

2 Binding relationships: Determining nodes bound by other network elements

Interaction networks were visualized with transcription factors represented as yellow squares and target genes as purple circles, facilitating the analysis of transcriptional regulation patterns.

TF-target gene interaction graph: This module displays the network displaying the number of genes targeted to regulate transcription factors using the data received from the server. The yellow square represents the transcription factor and the purple circle represents the target gene. The interaction between the targeted genes and TF can help in understanding the organization of transcriptional regulation and how TF modulate the expression of target genes (Kulkarni et al., 2018).

**Gene expression module:**

The expression values of target genes regulating TF were curated from RNA-seq. The expression of target genes are significantly influenced by the TF they regulate. The expression of target genes influences various biological processes. The normalized expression of the target genes in the network is displayed as a bubble chart, which was generated using the Plotly Python library. The x-axis represents the genes, and the y-axis represents the normalized expression values. The size of the bubble represents gene expression. The larger the bubble, the higher the expression of that gene.

**Network Ontology module:**

The Gene Ontology annotation information curated from the TAIR database is displayed on the Gene Ontology tab when the node is clicked. When a node is clicked, a request to fetch the data is sent from JavaScript using Ajax to the Flask file, which retrieves the information stored in MongoDB using the PyMongo library (https://pymongo.readthedocs.io/en/stable/), which connects MongoDB and Python, and displays the data in the form of a table in HTML. Three tables are displayed: Biological process (a process that is accomplished by multiple molecular activities), molecular function (gene activity at the molecular level), and cellular components (the location where molecular processes occur) (Thomas, 2016, 15–24).

The functional enrichment tab consists of the top 20 GO enriched terms plots, which were generated using g:Profiler REST API (an R package used for the functional enrichment analysis of genes) (Kolberg et al. 2020).

The pathway analysis tab displays the pathways for the genes involved in the network. It displays gene information, pathways associated with the gene, pathway image, and description. This table and image are displayed using the Biopython module (https://biopython.org/) (a tool for biological computation) which interacts with (Kyoto Encyclopedia of Genes and Genomes (API) and displays graphical representations of cellular processes (Minoru Kanehisa & Susumu Goto, 2000).

**Motif Visualization module:**

The gene structure was generated using the feature viewer library. The blue rectangle represents the promoter region and the gray rectangle represents the exon. The binding site is shown in red in the gene structure. The sequence information of the binding sites was used to create the logo displayed below in image form using the Logomaker Python package.

**Comparison module:**

To compare the gene regulatory network graph under different conditions, the two drop-downs are used to select the two conditions that the user wants to compare. Upon selecting the option from the drop-down, the data are split into two parts depending on the conditions selected: the network graph of the selected condition (generated through D3js), its network property (calculated using the NetworkX library), and the common modules (calculated using the greedy modularity communities library of NetworkX and displayed using the D3js library) in the form of a chart are displayed below the drop-down. Similarly, the width of the edge represents the weight or value with which node is associated with another node

### Results and discussion

#### Grad-CAM as an Explainable AI Framework for TFBS Identification Using Causal Networks

Understanding TFBS and their regulatory mechanisms is crucial for uncovering gene regulation processes. As deep learning models become increasingly sophisticated in detecting TFBS, the need for explainability grows. To address this challenge, we implemented Gradient Weighted Class Activation Mapping (Grad-CAM), a method initially introduced by Selvaraju et al. (2017) for visualizing deep learning decisions in medical imaging, to sequence data and TF causal networks. This adaptation of Grad-CAM aims to identify the contextual and gene features in TF causal networks that most significantly contribute to TF binding site discovery under specific conditions. By incorporating causal network information and sequence-level features, this approach enhances the interpretability of complex models and provides insights into the mechanisms governing TF binding. To compute the importance of each node in the TF causal network for identifying DNA binding, we first calculate the gradient of the output y (the likelihood of TF binding) with respect to the node-specific activations. This is expressed as $\frac{dy}{dh_i}$ represents the feature vector of node $v_i$ in the network. The importance weight $\alpha_i$ for each node$v_i$ is derived by averaging the gradient components over all dimensions $d$ of $h_i$:

$$\alpha_i = \frac{1}{d}\sum_{k=1}^{d} \frac{dy}{dh_i^{(k)}}$$

where $h_i^{(k)}$ is the *k*-th dimension of $h_i$. To compute the contribution of node $v_i$, we use the ReLU activation function to retain only positive contributions, ensuring that the negative gradients do not detract from the importance measure. The node score is given by:

$$NodeScoreforv_i = ReLU(\alpha_i . h_i) = max(0, \alpha_i . h_i)$$

The total contribution of the network is then calculated by summing the ReLU-activated scores across all nodes in the graph:

$$S_{Grad-CAM} = \sum_{v_i \in V} ReLU(\alpha_i . h_i)$$

This ensures the distribution of importance across all nodes reflects their relative roles in determining the TF binding under specific conditions.

In our implementation of Grad-CAM, we selected the normalization layer following the XGboost layer as our layer of interest **(Figure 3)**. This layer contains distribution maps for diverse nodes present in the causal network. Adhering to the weighting methodology proposed by Grad-CAM, we quantified the significance of these nodes and computed a weighted summation of all distribution maps. This aggregated map effectively highlights the nodes that are pivotal to binding activities. Subsequently, we conducted enrichment analysis to determine whether each unique k-mer was enriched among all k-mers with the highest average Grad-CAM scores. This analysis was performed across 35 TFs utilized in the study. We also conducted a comprehensive performance evaluation, comparing Grad-CAM-derived motifs with those reported by CTF-BIND and experimentally validated motifs from the JASPAR. This comparative analysis serves to validate the efficacy of Grad-CAM implementation in identifying biologically relevant sequence features and to assess its performance against established methodologies in the field of TFBS identification. Grad-CAM was extended to analyze sequence-level features by incorporating sequence motifs and causal network information. For each sequence, the model first computes a Grad-CAM score profile highlighting the contextual features most strongly associated with TF binding status. The relevance of sequence features, such as k-mers, was evaluated by identifying those with the highest average Grad-CAM scores. In the causal network, Grad-CAM was applied to identify key nodes and edges contributing to the performance ot the model. The network represents TFs and their target genes as nodes, with directed edges encoding causal relationships and regulatory influences. By analyzing the Grad-CAM score distributions across the network, nodes with significant contributions to the binding sites were identified.

**Case Study 1:**

To identify key regulatory nodes within the TF causal networks and understand their influence on TF binding, we applied Grad-CAM, visualizing the importance of each node (TF or gene) for the discovery of TF binding to specific DNA sequences under particular conditions. Applying Grad-CAM to the WRKY33 causal network pinpointed key structural and sequence features influencing its binding, with top-scoring k-mers exhibiting 84.94% concordance with known WRKY33 motifs in CTF-BIND. Five key nodes (WRKY25, MPK3, STZ, MKK6, and MYB51) exhibited high Grad-CAM scores (>0.8), **(Figure 9)** and their *in silico* removal from the network abolished accurate WRKY33 motif identification, demonstrating their crucial role in determining WRKY33 binding specificity. To validate these computational findings, we conducted a literature review and revealed strong experimental support for the identified interactions: WRKY33 is established as a condition-dependent master regulator in plant specialized metabolism, particularly in the camalexin and 4OH-ICN pathways, coordinating with MYB51 in *Arabidopsis thaliana* for gene expression and metabolic fluxes (Barco et al. 2020; Chen et al. 2024). Both WRKY33 and WRKY25, members of the WRKY family involved in stress responses, exhibit distinct regulatory roles in *Gossypium hirsutum*, with WRKY33 acting as a stronger negative stress regulator (Ehsan et al. 2023; Li et al. 2011). While direct evidence for WRKY33 interactions with MPK3, STZ, and MKK6 was limited, but their known roles in stress response pathways (MPK3 in MAP kinase signaling, STZ in salt tolerance, and MKK6 also in MAP kinase cascades interacting with WRKY TFs) provide compelling contextual support (Andreasson et al. 2005; Mao et al. 2011; Zhou et al. 2020; Han et al. 2019). This integrated approach of computational analysis and experimental validation highlights the complex regulatory mechanisms governing TF binding, demonstrating that WRKY33's binding is influenced not only by its core motif but also by interactions within its regulatory network.

**CTF-BIND database application demonstration**

This case study demonstrates the utility of CTF-BIND-DB in dissecting the dynamic transcriptional regulatory network of HSFA1 (Heat Shock Transcription Factor A1) under heat stress in *Arabidopsis thaliana.* Heat shock-responsive genes, encoding proteins like heat shock proteins (HSPs), are crucial for maintaining cellular homeostasis under heat stress by preventing protein denaturation and aggregation (Liu et al., 2011; Ohama et al., 2017). Using CTF-BIND-DB, we investigated how HSFA1's interactions shift dynamically across different heat stress time points (**Figure 11)**.

**Network module:** We began by exploring the Network module of CTF-BIND-DB, focusing on HSFA1's interactions under heat stress across seven time points. It revealed that HSFA1 targets 47 heat shock-responsive genes, including key HSPs like HSP70, HSP90, and HSP101, as well as several small HSPs (sHSPs). HSFA1 exhibited high degree centrality (0.85) and closeness centrality (0.76), confirming its central role as a hub within the heat stress network. The betweenness centrality was also high (0.71), further suggesting that HSFA1 acts as a key information conduit within the network. High centrality measures indicate HSFA1's importance in coordinating the heat stress response. Its numerous connections and strategic position within the network allow it to rapidly influence the expression of a large number of downstream genes. A dynamic force-directed graph visualized the network, demonstrating that several key target genes clustered in modules enriched for heat-responsive pathways. The network's clustering coefficient of 0.34 suggests a moderately interconnected structure, allowing for both local and global communication within the network.

**Interaction Module:** The TF-target gene interaction graph within the Interaction module visually confirmed the direct interactions between HSFA1 and its 47 target genes. Analysis of "Bound by relationships" indicated that HSFA1 is tightly bound to the promoter regions of these genes, particularly during the initial stages of heat stress (0-3 hours). The "Targeting relationships" table revealed that HSP70 exhibited the strongest targeting relationship with HSFA1, with an edge weight of 0.92, suggesting its crucial role of this chaperone protein in mitigating the immediate effects of heat stress. The temporal binding pattern suggests a rapid and direct transcriptional activation mechanism.

**Gene Expression Module:** Analysis of normalized expression data across seven time points (0.5, 1, 3, 6, 12 and 24 hours) using the Gene Expression module revealed a coordinated expression pattern. HSFA1 expression peaked within the first 3 hours, followed by a gradual decline after 6 hours. Target genes such as HSP70 and HSP101 followed a similar trend, showing a significant expression increase during early stress stages. The bubble chart further emphasized the high expression levels of HSP70 and sHSPs under heat stress. The transient expression pattern of HSFA1 and its targets reflects the dynamic nature of the heat stress response. The rapid induction of HSPs at early time points allows the plant to quickly counteract the initial effects of heat stress, while the subsequent decline in expression prevents unnecessary energy expenditure.

**Network Ontology Module:** To understand the biological functions of HSFA1's target genes, we performed gene set enrichment analysis using the Network Ontology module, revealed significant enrichment of terms related to "Response to heat" (GO:0009408, FDR = 0.0237), "Cellular response to heat" (GO:0034605, FDR = 4.45E-06), "Response to hydrogen peroxide activity" (GO:0042542, FDR = 0.00875), and "Cellular response to hypoxia" (GO:0071456, FDR = 0.751). KEGG pathway analysis showed enrichment of "Protein processing in the endoplasmic reticulum" (FDR = 1.33e-12) and "Regulation of HSF1-mediated heat shock response" (FDR = 0.0124). The enrichment of these GO terms and KEGG pathways highlights the critical role of protein quality control, including protein folding, trafficking, and degradation, in the heat stress response. The involvement of "Response to hydrogen peroxide activity" suggests a crosstalk between heat stress and oxidative stress signaling.

**Motif Visualization Module:** Promoter analysis using the Motif Visualization module identified highly conserved heat shock elements (HSEs) in the promoter regions of target genes. HSP70, for instance, had three binding sites for HSFA1, with a binding affinity score of 0.87. The motif logo visually represented the sequence conservation of HSE motifs. The presence of HSEs directly confirms the mechanism of HSFA1-mediated transcriptional regulation, demonstrating its direct binding to target gene promoters.

**Comparison Module**: Comparing the HSFA1 network under heat stress with that under cold stress revealed significant differences. Under heat stress, HSFA1 targeted 47 genes, whereas only 12 overlapping genes were targeted under cold stress. Heat-specific genes like HSP70 and HSP101 showed negligible expression under cold stress. Network density was higher under heat stress (0.43) than under cold stress (0.21). This comparison demonstrates the specificity of HSFA1's regulatory role in the heat stress response. The distinct target gene sets and network properties under different stresses highlight the fine-tuned control of gene expression in response to specific environmental cues.

**Case Study 2: Deciphering Salt-Tolerance regulatory Mechanisms**

Salinity stress, a major abiotic stressor, disrupts ion homeostasis and osmotic balance in plants, significantly impacting growth and productivity (Joshi et al., 2022). This case study utilizes CTF-BIND-DB to explore the regulatory networks of key salt-responsive TFs, SOS1 (Salt Overly Sensitive 1) and WRKY8, providing insights into the mechanisms of salt tolerance (**Figure 11)**.

**Network Module:** Analysis using the Network module revealed that WRKY8 and SOS1 targeted 54 salinity-responsive genes, including those involved in ion transport (e.g., NHX1) and osmotic regulation (e.g., P5CS1). WRKY8 exhibited high degree centrality (0.88) and betweenness centrality (0.79), highlighting its central role in the salinity stress network. A force-directed graph visualized clusters of genes enriched in pathways like Na+/K+ ion transport and ABA signaling. The network's clustering coefficient was 0.42, indicating a well-connected structure facilitating efficient communication between network components. The high centrality measures of WRKY8 underscore its importance as a major regulator of the salt stress response. The clustering of genes involved in ion transport and ABA signaling suggests coordinated regulation of these crucial processes.

**Interaction Module:** The Interaction module provided detailed information on the interactions between SOS1, WRKY8, and their target genes. The TF-target gene interaction graph visualized these connections, revealing that SOS1 primarily targeted genes involved in ion transport and homeostasis, while WRKY8 targeted a broader set of genes involved in various stress response pathways, including ABA signaling and antioxidant defense. This functional divergence suggests specialized roles for these TFs. SOS1 appears to be directly involved in maintaining ion balance, a primary response to salt stress, while WRKY8 coordinates a more diverse set of downstream responses, including hormonal signaling and antioxidant defense, to mitigate secondary stress effects.

**Gene Expression Module:** Analysis using the Gene Expression module revealed distinct temporal expression patterns for SOS1 and WRKY8 across four salinity stress time points (0, 6, 12, and 24 hours). Both WRKY8 and SOS1 exhibited peak expression at 6 hours, coinciding with the induction of NHX1 and other ion transporters. Expression levels gradually declined by 24 hours, suggesting an early response mechanism. Bubble charts by the module illustrated co-expression clusters of genes involved in ion transport and osmotic adjustment. The coordinated peak expression of SOS1 and WRKY8 at 6 hours suggests a synchronized activation of salt tolerance mechanisms. The subsequent decline in expression indicates the transient nature of the initial response, potentially followed by other regulatory mechanisms for long-term adaptation.

**Network Ontology Module:** Network Ontology analysis revealed significant enrichment of GO terms related to "Response to salt stress" (GO:0009651, FDR = 0.017), "Ion transport" (GO:0006811, FDR = 0.009), and "Osmotic regulation" (GO:0006970, FDR = 0.023). KEGG pathway analysis showed significant enrichment for "ABC transporters" (K02000, FDR = 1.22e-11) and "Proline biosynthesis" (K00931, FDR = 4.34e-05). The enrichment of these GO terms and KEGG pathways confirms the involvement of the identified genes in key salt tolerance processes, such as maintaining ion homeostasis, osmotic adjustment, and detoxification. The enrichment of "ABC transporters" highlights the importance of active transport mechanisms in coping with salt stress, as earlier seen in some studies (Dahuja et al., 2021), while "Proline biosynthesis" indicates the role of compatible solutes in osmotic adjustment.

**Motif Visualization Module:** Motif analysis using the Motif Visualization module identified conserved W-box elements (T(T/A)GAC(C/T)) in the promoter regions of WRKY8 target genes, confirming its direct DNA binding. SOS1 promoters displayed enriched binding sites for HSEs and ABREs, suggesting potential crosstalk with heat stress and ABA signaling. Promoters of NHX1 and P5CS1 had high binding affinity scores of 0.89 and 0.85, respectively, suggest strong regulatory control of these key salt tolerance genes. The presence of W-boxes confirms WRKY8's direct transcriptional regulation of its target genes. The identification of HSEs and ABREs in SOS1 promoters suggests potential integration of salt stress responses with other stress signaling pathways.

**Comparison Module:** Comparing WRKY8's regulatory network across the four time points (0, 6, 12, and 24 hours) using the Comparison module revealed dynamic changes in network structure. At 6 hours, WRKY8 targeted 54 genes, while this number dropped to 21 at 24 hours. Cross-condition analysis revealed minimal overlap between salinity and drought stress networks. The decrease in WRKY8 target genes at later time points suggests a shift in regulatory priorities as the plant adapts to prolonged salt stress. The minimal overlap between salinity and drought stress networks highlights the specificity of transcriptional responses to different abiotic stresses.

These case study exemplifies the power of CTF-BIND-DB in dissecting complex regulatory networks under abiotic stress. By integrating diverse data types and providing user-friendly modules, CTF-BIND-DB facilitates a deeper understanding of TF function and stress response mechanisms. These findings provide valuable insights into plant stress biology and pave the way for developing strategies to enhance crop resilience to environmental challenges.
